# Deep Learning and Association Rule Mining for Predicting Drug Response in Cancer. A Personalised Medicine Approach

**DOI:** 10.1101/070490

**Authors:** Konstantinos Vougas, Magdalena Krochmal, Thomas Jackson, Alexander Polyzos, Archimides Aggelopoulos, Ioannis S. Pateras, Michael Liontos, Anastasia Varvarigou, Elizabeth O. Johnson, Vassilis Georgoulias, Antonia Vlahou, Paul Townsend, Dimitris Thanos, Jiri Bartek, Vassilis G. Gorgoulis

**Author notes:** To whom correspondence should be addressed: Vassilis G. Gorgoulis; Jiri Bartek.

## Abstract

A major challenge in cancer treatment is predicting the clinical response to anti-cancer drugs for each individual patient. For complex diseases such as cancer, characterized by high inter-patient variance, the implementation of precision medicine approaches is dependent upon understanding the pathological processes at the molecular level. While the “omics” era provides unique opportunities to dissect the molecular features of diseases, the ability to utilize it in targeted therapeutic efforts is hindered by both the massive size and diverse nature of the “omics” data. Recent advances with Deep Learning Neural Networks (DLNNs), suggests that DLNN could be trained on large data sets to efficiently predict therapeutic responses in cancer treatment. We present the application of Association Rule Mining combined with DLNNs for the analysis of high-throughput molecular profiles of 1001 cancer cell lines, in order to extract cancer-specific signatures in the form of easily interpretable rules and use these rules as input to predict pharmacological responses to a large number of anti-cancer drugs. The proposed algorithm outperformed Random Forests (RF) and Bayesian Multitask Multiple Kernel Learning (BMMKL) classification which currently represent the state-of-the-art in drug-response prediction. Moreover, the *in silico* pipeline presented, introduces a novel strategy for identifying potential therapeutic targets, as well as possible drug combinations with high therapeutic potential. For the first time, we demonstrate that DLNNs trained on a large pharmacogenomics data-set can effectively predict the therapeutic response of specific drugs in different cancer types. These findings serve as a proof of concept for the application of DLNNs to predict therapeutic responsiveness, a milestone in precision medicine.

## INTRODUCTION

Predicting the clinical response to therapeutic agents is a major challenge in cancer treatment. Ultimately, the ability to generate genomic-informed, personalized treatment with high efficacy, is dependent upon identifying molecular disease signatures and matching them with the most effective therapeutic interventions. The advent of multiple platforms providing “omics” data permits scientists to dissect the molecular events that are known to drive carcinogenesis^1^ and alter major downstream processes, such as gene expression^2^. Nonetheless, effective translation of the growing wealth of high-throughput profiling data into a personalised treatment strategy required by precision medicine, has been daunting and without noteworthy success^3^.

The successful identification of effective anti-cancer drugs has been primarily hindered by the lack of reliable preclinical models. Although individual cancer cell lines lack the complexities of clinical cancer tissues^4^, when compiled in large panels, have been reported to recapitulate the genomic diversity of human cancers^5^. Such panels can be readily utilised as platforms upon which expert systems for the prediction of pharmacological response may be developed. To facilitate such a task three large-scale cell panels containing pharmacogenomics data have been made available to the public domain: **a)** the “Cancer Cell Line Encyclopedia” (CCLE)^6^, **b)** the Genomics of Drug Sensitivity in Cancer (GDSC)^7^ and **c)** the NCI-60^8^. To identify predictive biomarkers, these consortia have analysed the molecular profiles of over 1000 cancer cell lines and drug profiles of a large number of anti-cancer drugs.

The availability of these large data sets of cell-line panels along with the availability of new computational technologies has propelled a widespread effort to perform parallel analyses across cell lines, in order to extract novel information and define predictive biomarkers. However, while large data sets of pharmacogenomic profiles have been compiled with detailed molecular features and drug responsiveness, well-validated computational approaches defining biologically relevant rules, and algorithms able to accurately predict the responsiveness to a specific therapeutic drug are lacking. Although data mining algorithms are supposed to analyse large volumes of data and uncover hidden relationships of potential clinical significance, today’s complex “omics” datasets have appeared too multi-dimensional to be effectively managed by classical Machine Learning algorithms^9^. Deep Learning neural networks (DLNN), on the other hand, have the ability to “understand” complexity and multidimensionality and have been effectively applied in various fields (e.g. image analysis, text mining, etc.) with increased classification accuracy compared to classical computation methods^10^. DLNN is based on the modelling of high-level neural networks in flexible, multilayer systems of connected and interacting neurons, which perform numerous data abstractions and transformations^11^. In a recent surge of interest, DLNN has been effectively applied to extract features from various large and complex data sets, including predicting drug-target interactions^12^, drug toxicity in the liver^13^ and pharmacological properties of drugs^14^, among others. Altogether, studies using the DLNN architecture demonstrate its suitability for the analysis of complex biological data, as it can automatically construct complex features and allows for multi-task learning^15^.

We designed a bioinformatics pipeline for handling multiple layers of molecular profiling information extracted from publicly available pharmacogenomics resources, in order to produce an expert system that, with demonstrated efficiency, could predict pharmacological responses to a large number of drugs over a broad panel of cancer cell-lines (**Figure 1**). Specifically, we performed feature selection in the form of association rules and utilized the selected features to train the state-of-the-art DLNN to predict pharmacological response in a blind set. The association rules are treated as a novel meta-dataset which is utilised in the form of proofs of concept for knowledge extraction biomarker discovery and novel therapeutic target identifications. To our knowledge, this is the first time that the DLNN framework is systematically applied to predict drug efficacy in cancer treatment.

**Figure 1.**
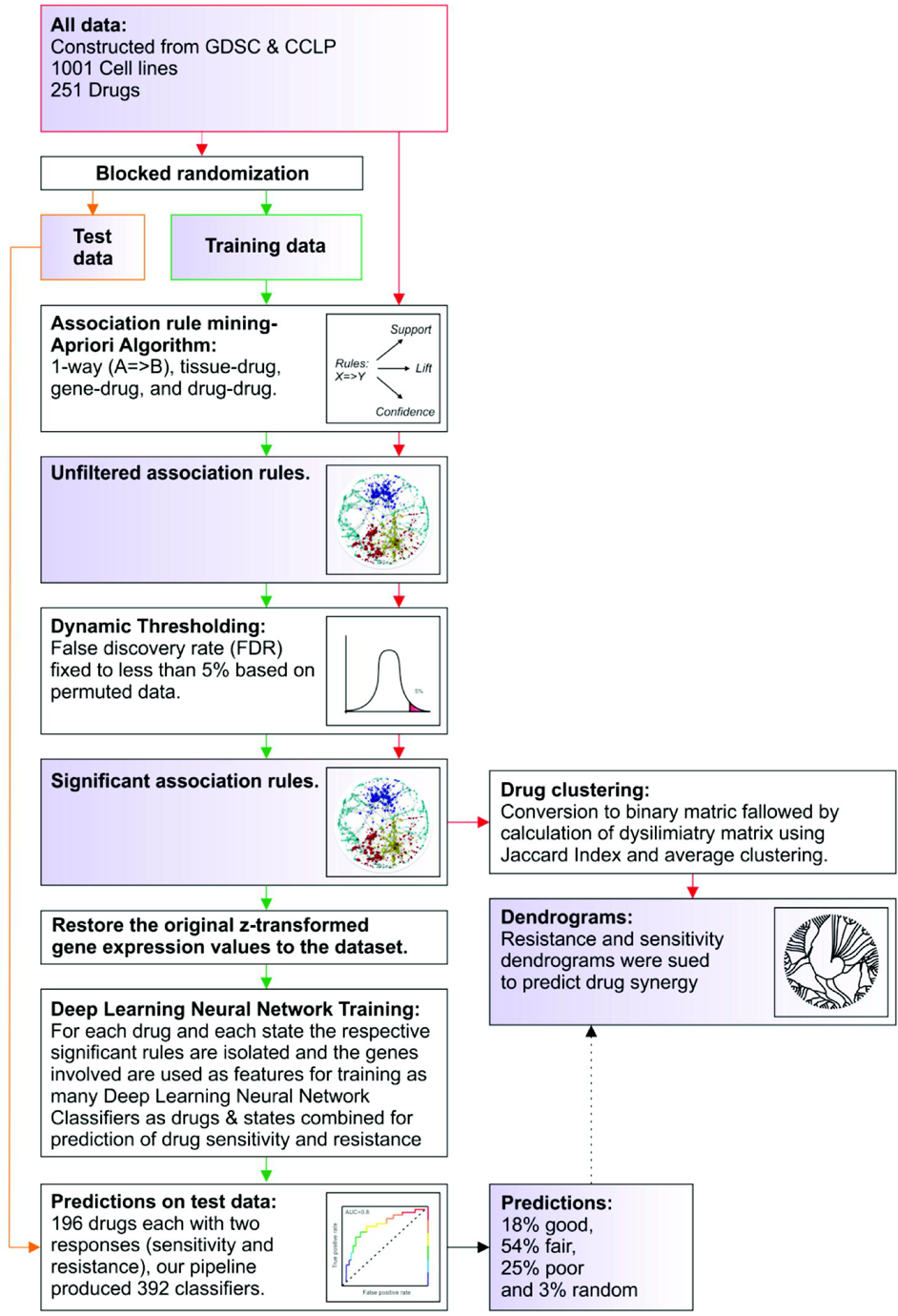
Schematic representation of the study design and bioinformatics pipeline. The full data set was constructed using the GDSC and CCLP databases. Test data and training data sets were created using blocked randomisation. The use and progress of each data set is depicted by coloured arrows; full data set (red), test set (orange) and training set (green). The training and test set were used for deep learning and subsequent predicting of sensitivity and resistance. The full data set was used for clustering and dug synergy predictions. Association rule mining was used for feature selection (from >60,000 features) of pharmacogenetics data. Significant association rules were defined by dynamic thresh-holding. Rules derived from the training data set were used for deep learning. DLNN classifiers were applied for prediction of drug sensitivity and resistance. Extracted information was validated using the test set. Genes from rules derived from the full data set were used to construct a dissimilarity matrix based on the Jaccard index. Drugs were then clustered and predictions for drug synergy made.

## RESULTS

### 1. Dataset compilation

To initiate a bioinformatics pipeline for prediction of drug response based on molecular profiles of multiple cancer cell types, we generated a large-scale pharmacogenomics dataset for 1001 cancer cell lines and 251 anti-cancer drugs (**Figure 2, Methods – Datasets**). The new pharmacogenomics compilation was achieved by merging data from the Cosmic Cell-line project (CCLP) and GDSC. We used GDSC^7^ as our drug response data source for 251 therapeutic compounds, which provided IC-50 values for each compound, as well as information on tissue origin. Information on total gene mRNA expression, number of DNA copies and mutational status was obtained from the Cosmic Cell-line project (CCLP)^16^. CCLP was preferred as a data source since it provides profiles on 1,074 cancer cell lines and is not limited to the mutational status of only 1600 genes, as is the case with CCLE. Although NCI-60 contains the largest number of therapeutic compounds tested for pharmacologic activity, it was excluded as a data source, as the number of cell-lines presented is very low compared to the other resources used. This diminishes the effectiveness of NCI-60 to serve as a preclinical platform that can, at least partially, simulate clinically relevant tumour complexity.

**Figure 2:**
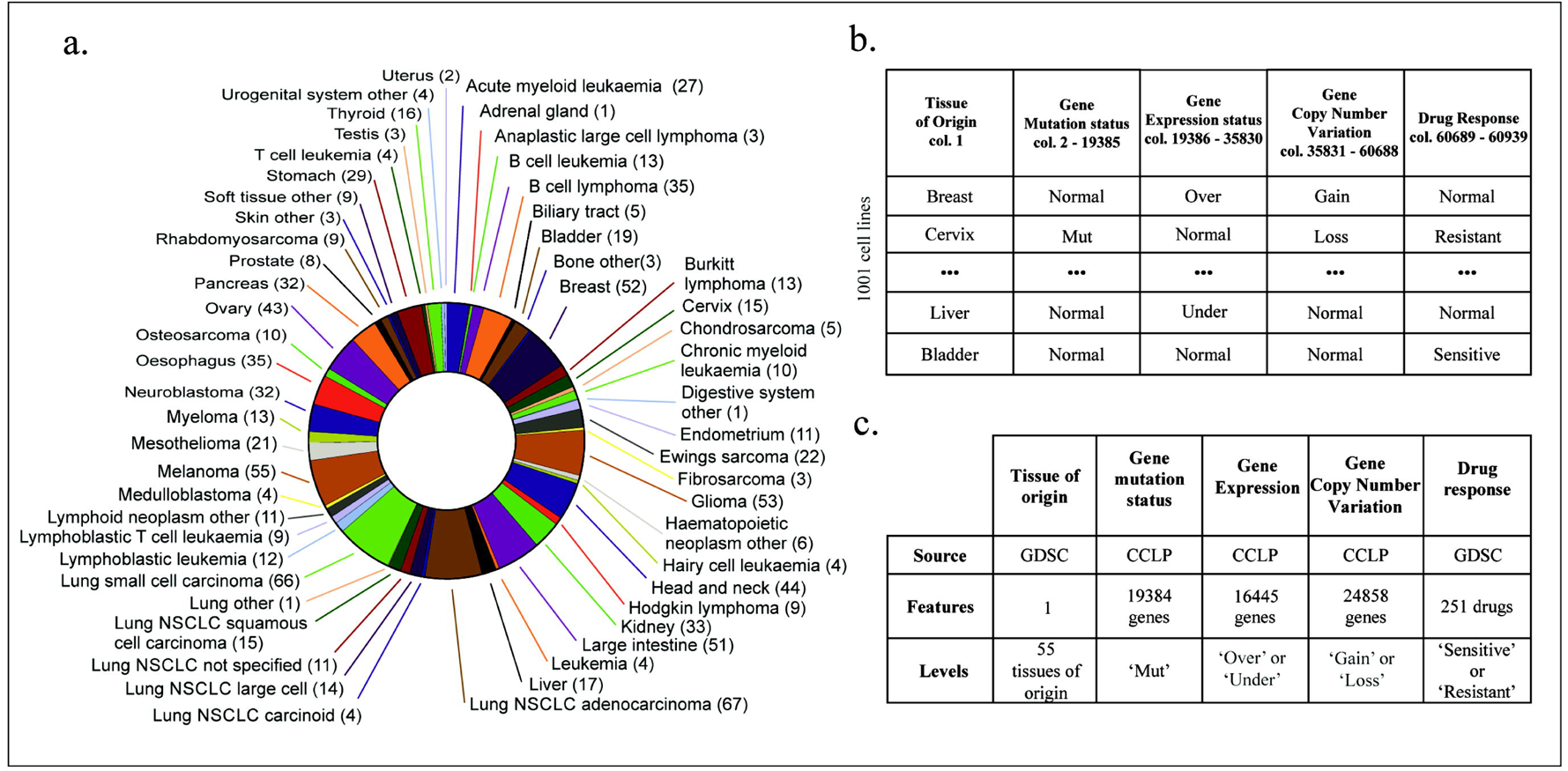
Description of full data set and summary of main data matrix. **a)** Tissue of origin of the 1001 cell-lines of the data-set**. b)** Summary of the main data matrix containing tissue of origin, mutation status, gene expression, copy number variation and drug response information for the 1001 cancer cell-lines. The actual matrix is available as an R data object (MASTER_MATRIX.RData) that can be accessed from the data folder located in the GitHub repository (**Methods – Data Availability**). **c)** Description of each data type used including source, number of features and levels.

### 2. Association Rule Mining

Given their molecular profiling data, both large cell-line panels (CCLE and GDSC) have been utilized in attempts to identify biomarkers for predicting drug response of specific cancer cell-lines^6, 7^. Previous efforts to define biomarkers of drug response primarily utilize elastic net regression, a penalized linear modelling technique, to identify cooperative interactions among multiple genes and transcripts across the genome and define response signatures for each drug^16^. While efficient, this algorithm suffers certain limitations since when used for feature selection, as described in previous studies^6, 7^, the derived results are simple associations between a single gene and drug response. If, however, one wishes to explore the relevance of a more complex feature-space relationship (two or three-way interactions among simple features in all possible combinations) to the drug response, the process is convoluted. This is primarily due to the fact that these algorithms fall-short in automatically evaluating all possible combinations including multi-way interactions of a large number of features against a response variable without further implementation. Furthermore, multi-feature models generated by such algorithms are difficult to interpret in terms of biological relevance. When utilised as a classifier to predict whether a sample will be resistant or sensitive to a drug, given its molecular profile, the elastic net algorithm does not perform optimally. This is due to the fact that at the core of the elastic net algorithm lays linear regression, as opposed to non-linear classifiers, such as Random-Forests and Kernel-based models. The later have been shown to outperform the elastic net algorithm in the task of actually predicting drug response, as demonstrated in a recent proof of concept study on a panel of 53 breast cancer cell lines evaluated for pharmacological response against 28 anti-cancer drugs^17^.

#### 2a. Apriori Algorithm

To overcome the primary limitations of the elastic net algorithm for feature selection, we applied a method used by large businesses to analyse the enormous volume of transaction data and discover all possible associations between the data features, namely Market Basket Analysis or Association Rule Mining. Previous studies moved along the same lines to produce easily interpretable logical rules out of similar pharmacogenomic datasets^5, 18^. The reason we selected to proceed with association rule mining was the fact that it provides an efficient big-data ready framework that is able to evaluate a huge sample space of associations among features including multi-way interactions with more than 30 different objective measures^19^. Additionally, the output of the algorithm comes in the form of easily interpretable rules, making knowledge extraction and meta-analysis a more straightforward process. Specifically, we applied the Apriori algorithm^20^ to extract significant associations from all of the possible combinations of the features from the main dataset (tissue of origin, gene expression, mutation status, CNV plus drug response), in order to generate a large rule-set, containing all tissue-to-gene, tissue-to-drug, gene-to-gene, gene-to-drug and drug-to-drug associations. The main bottleneck in the application of association rule mining in this study is the computationally intensive requirements. While this will likely improve as computing power increases, due to hardware limitations we maintained only the tissue-to-drug, gene-to-drug and drug-to-drug associations for the present study. Gene-to-gene associations, which constitute an enormous RAM intensive rule-set, were discarded. Details and metrics of the Apriori algorithm can be found in the **Methods - Association Rule Mining – Apriori Algorithm**. Relationships between confidence and support metrics (for top 10,000 1-way and 100,000 2-way rules) are visualized in the scatterplots in **Supplementary Figure 1.**

#### 2b. Dynamic Thresholding - Separating true rules from the noise

We devised a procedure that we named Dynamic Thresholding in order to select significant non-random rules by controlling the false discovery rate (FDR) to less than 5%. Dynamic Thresholding is based on running the Apriori algorithm on a permuted version of our initial dataset (**refer to Methods -Association Rule Mining – Apriori Algorithm / Dynamic Thresholding**). The biological relevance of the rules generated was examined in separate proofs of concept, as we show below.

### 3. Rule Verification

#### 3a. Proofs of concept

To validate the biological relevance of our statistically significant association rules, we examined whether known predictors of drug response are present in our rule set and whether drugs for a given target are present in sensitivity-associated rules along with the given target if mutated or over-expressed.

##### Proof of concept 1

We demonstrate that over-expressed NAD(P)H dehydrogenase 1 (NOQ1) and MDM2, a p53 inhibitor, which are known predictors of sensitivity for the drugs 17-AAG (Tanespimycin) and Nutlin-3, respectively^21, 22^, are present in our rule-set (**Supplementary Table 1 - 1-way rules**). Additionally, our rules indicate that EGFR over-expression and suppression are significantly associated with Lapatinib sensitivity and resistance respectively which is in agreement with previous findings which describe that EGFR expression can efficiently model lapatinib response ^23^(**Supplementary Table 1 - 1-way rules)**. Finally the *ABCB1* gene that encodes the Multidrug-Resistance-1 (MDR1) protein, was found in our rule set to be linked with resistance to multiple drugs when it is over-expressed (55 out of 57 drugs), while when suppressed it is linked with sensitivity (7 out of 9 drugs) (**Supplementary Table 1 - 1-way rules).**

##### Proof of concept 2

We performed two k-mean clustering schemes (**Methods - Association Rule Mining – Apriori Algorithm**) of the 1000 rules with the largest support (k=50 which represents the top 5% of the most frequently occurring molecular events related to the given condition) for the sensitivity response-state of drugs associated with (a) the ERK/MAPK signalling and (b) the PI3K signalling (**Supplementary Table 1 - 1-way rules, Supplementary Table 2**). The first clustering scheme revealed that the mutated BRAF which is central to ERK / MAPK signalling was present among the top 50 cluster centres (**Figure 3a**). Additionally, the ERK / MAPK -clustering revealed that the melanoma cell-lines were highly sensitive to BRAF and MEK inhibitors, a “prediction”, which can be verified in the literature with studies showing that combined BRAF and MEK inhibition is in fact, one of the most effective treatments for melanomas^24^ (**Figure 3a**). The actual **half maximal inhibitory concentration** (IC_50_) values of the drugs included in this group indicate increased sensitivity for melanoma cell-lines and for cell-lines carrying mutated BRAF as compared to the total dataset (*p*-value <0.05) (**Figure 3b**). The second clustering scheme revealed the presence of the mutated PTEN among the top 50 cluster centres (**Figure 3c**). PTEN is a direct PIK3CA suppressor^25^ that is frequently mutated in cancer with loss-of-function mutations,^26^ which in turn leads to increased PIK3CA activity. Notably, mutated PIK3CA was also present in the mutated-PTEN cluster (**Figure 3d**). Given that both, PTEN and PIK3CA, belong to the same pathway, the fact that the onco-suppressor (PTEN) is deactivated at the same time that the oncogene (PIK3CA) is further activated by hot-spot gain-of-function mutations could be visualised as a variation of the Knudson double-hit hypothesis^27^. The aforementioned facts confirm that rule-derived clustering schemes, as the ones currently described, provide relevant insights regarding the molecules that are related to responsiveness to certain drug-classes. After demonstrating that rule-clustering delivers relevant results, we present an example of how the rules can be used to gain novel insights on biomarker discovery for drug response. The PI3K-clustering, links the suppression of the ID1 gene to sensitivity to 10 out of 16 drugs targeting the PI3K pathway with high lift and support values (**Figure 3c**). Inhibitor of DNA binding 1 (ID1) is a transcription regulator, widely reported as linked to tumour metastasis when over-expressed^28,29^ and known to activate the PI3K pathway^30^, while inhibition of ID1 expression suppresses cancer invasion and progression^31, 32^. Based on the TCGA gene expression data, which are derived from analysing patients and therefore constitute a direct link of our findings with clinical settings (**Supplementary Table 3**), we noticed a high percentage of ID1 suppression in several cancer types (e.g. Breast invasive carcinoma [TCGA code: BRCA], Lymphoid Neoplasm Diffuse Large B-cell Lymphoma [TCGA code: DLBC], etc.) (**Figure 3f**). IC_50_ heatmaps (**Figure 3e**) indicate that cell lines under-expressing ID1 are much more responsive to inhibitors targeting the PI3K pathway in contrast to cell lines over-expressing ID1 which seem to be resistant towards the same inhibitors (p<0.01). These results indicate that apart from being used as a therapeutic target *per se*, ID1 could be utilised as a biomarker for responsiveness to PI3K-targeted therapies, as its expression seems to distinguish sensitive from resistant cell lines more efficiently than the actual PIK3CA mutation status (**Figure 3e**). In agreement with ID1 expression data from TCGA (**Figure 3f)**, PI3K/AKT/mTOR inhibitors have been proven beneficial for the treatment of Acute Myeloid Leukemia [TCGA code: LAML]^33^ which shows low ID1 gene-expression levels in more than 90% of the recorded clinical cases. Additionally, Bladder Urothelial Carcinoma [TCGA code: BLCA] which demonstrates high ID1 levels in approximately 60% of the recorded clinical cases, respond poorly to PI3K mono-therapy^34^. Within this context, we recently depicted that chronic expression of the tumor-suppressor p21^WAF/Cip^^1^, in a p53-deficient environment, unrevealed an oncogenic behaviour generating aggressive and chemo-resistant clones. Surprisingly, and in line with the above observations ID1 was found up-regulated in these cells^2^. Moreover, in the ID1 rule-cluster, over-expression of 4 other genes was found to be highly related with sensitivity to PI3K-pathway inhibitors, namely *ZNF22*, *GMIP*, *LYL1* and *SAMSN1* (**Figure 3d)**. Interestingly, LYL1 (Lymphoblastic Leukemia Associated Hematopoiesis Regulator 1) is known to be implicated in the development of leukemias^35^ and lymphomas^36^, both representing promising target groups for anti-PI3K/mTOR agents^37, 38^. SAMSN1 (SAM Domain, SH3 Domain And Nuclear Localization Signals 1) is an intriguing case since it appears to act as a tumour suppressor in certain malignancies such as multiple myeloma^39^, gastric cancer^40^, lung cancer^41^ and hepatocellular carcinoma^42^, whereas its over-expression has been associated with poor survival in glioblastoma multiforme^43^; a malignancy where drug resistance represents a major obstacle^44^. Its detection in the rule-set concurs with recent developments suggesting that targeting the PI3K pathway could be a potential therapeutic option to overcome drug resistance in glioblastoma multiforme^45^.

**Figure 3:**
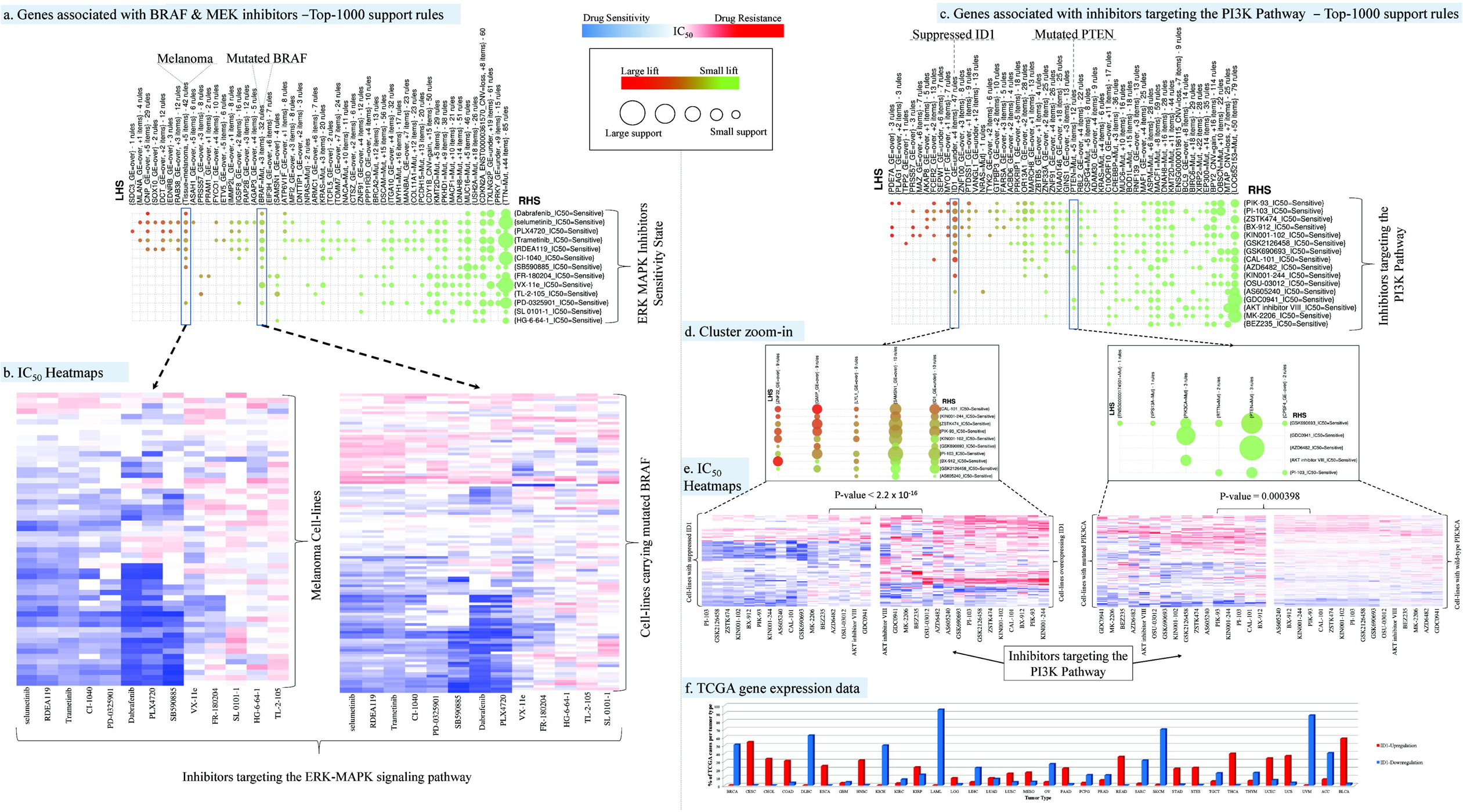
Association Rules visualization incorporating analysis and insights on the data. **a)** Group-wise Association Rules visualization by k-means clustering k=50 of the 1000 one-way rules with the largest support, for the sensitivity state of drugs targeting the ERK-MAPK signalling pathway. **b)** IC_50_ heatmaps of drugs targeting the ERK-MAPK signalling pathway for melanoma cell-lines and for cell-lines carrying mutated BRAF. **c)** Group-wise Association Rules visualization by k-means clustering k=50 of the 1000 one-way rules with the largest support, for the sensitivity state of drugs targeting the PI3K signalling pathway. **d)** Zoom-ins of the ID1 and PTEN clusters presented in section-c. **e)** IC_50_ heatmaps of drugs targeting the PI3K signalling pathway for cell-lines over and under-expressing ID1 and for cell-lines carrying wild-type and mutated PIK3CA. **f)** Percentage of TCGA cases per tumour type with ID1 levels one standard deviation above and below the mean. P-values were derived by the non-parametric two-tailed Wilcoxon test.

##### Proof of concept 3

The following two proofs of concept indicate how the association rules, when allowing for interactions (2-way), can be used to gain further insight in the molecular mechanisms of drug resistance in Small-cell lung cancer (SCLC) and identify potential points of intervention.

##### Proof of concept 3a

The 1-way rules, indicate a large pattern of multi-drug resistance (93 drugs) concerning SCLC (**Supplementary Table 1 -1-way rules**). SCLC accounts for approximately 15% of all lung cancers^46^ and is considered one of the most aggressive forms of lung cancer mainly due to rapid development of multi-drug resistance^47^ which is in agreement with our finding. The 2-way rules (**Supplementary Table 1 - 2-way rules**), indicate that the Growth hormone releasing hormone (GHRH) over-expression greatly increases the lift-value (hence statistical significance) to 39 of the above drugs. It is known that inhibiting GHRH activity using antagonists wields high anti-tumour activity by impending cell proliferation^48^. Furthermore, GHRH activity has been linked to drug-resistance in triple negative breast cancer^49^. We hereby demonstrate that by including interactions in association rule mining, we are able to infer in this particular proof of concept that GHRH antagonists could be potentially used in combination therapy schemes with specific chemotherapeutic agents for the effective treatment of SCLC, which is further supported by the fact that synergistic action of GHRH antagonists with Docetaxel has been successfully demonstrated for the treatment of non-small cell lung cancer^50^.

##### Proof of concept 3b

With the 1-way rules (**Supplementary Table 1 - 1-way rules**), we observe statistically significant resistance to Obatoclax-Mesylate, a BCL-family inhibitor, with a lift-value of 2.47 in 22 out of 66 SCLC cell-lines (33.3%). With the 2-way rules (**Supplementary Table 1 - 2-way rules**), we note that SMAD3 down-regulation greatly increases the lift-value to 4.77, since resistance to Obatoclax-Mesylate is observed in 9 out of 14 SCLC cell-lines under-expressing SMAD3 (64.3%). SMAD3 is known to promote apoptosis through transcriptional inhibition of BCL-2^51^. SCLC cell lines under-expressing SMAD3 clearly possess increased levels of BCL-2, which correlates well with the phenotype of resistance to a BCL-2 inhibitor, such as Obatoclax-Mesylate. In this example, association rule mining precisely elucidated a specific part of the resistance mechanism of SCLC to BCL-family inhibitors, by highlighting a unique molecule that presents high mechanistic relevance to BCL-inhibition.

##### Proof of concept 4 (Experimental)

To provide a proof of concept for the predictive ability of the association rule mining algorithm at the experimental level, we utilised our rule-set to identify potential candidate genes that upon silencing should affect the drug resistant phenotype by conferring increased sensitivity to a specifically applied treatment, namely Doxorubicin. Selection of the genes for experimental interrogation was based on an algorithm implementing the following strategy:

*i*) Over-expression of the candidate gene should be significantly connected to Doxorubicin resistance through a rule attaining a minimum 33% confidence value, which was set empirically and is indicative of the association power between the gene over-expression and the drug resistance phenotype. All the resulting rules where sorted (prioritized for consideration) from largest to smaller lift values.

*ii*) Over-expression of the specific gene should be connected through significant rules to as many (other than Doxorubicin) drug resistance phenotypes as possible.

*iii*) Suppression of the specific gene should be connected through significant rules to as many, other than Doxorubicin, drug sensitivity phenotypes as possible.

*iv*) Optionally, gene functionality is assessed for its biological relevance in the overall concept of drug resistance by utilizing prior knowledge.

The gene targets relevant to Doxorubicin resistance were “data-mined” from our rule-set and were compiled in **Supplementary Table 4 – “Targets selection”,** with all the metrics required by the aforementioned algorithm. We picked three genes from the list of potential targets for experimental validation. These genes were:

a. *Membrane Associated Guanylate Kinase, WW And PDZ Domain Containing 3* (*MAGI3*),
b. *Premature Ovarian Failure 1B* (*POF1B*), and
c. *Protein Disulfide Isomerase Family A Member 3* (*PDIA3*).

Validation of gene targets under the current scheme is meant as a proof-of-concept; hence a thorough validation of the targets is out of the scope of the current manuscript. We therefore decided to select three targets from the list which ideally should span the list from top to bottom, thus better illustrating its potential and at the same time maintain high metrics [based on strategy rules (*ii*) and (*iii*)] and present interesting functional properties.

MAGI3 was selected as it was found within the top-5 targets presenting the highest lift values for their association with Doxorubicin resistance and is the second best regarding strategy rules (*ii*) and (*iii*) (**Supplementary Table 4 - “Targets selection”, column:” Confirmatory_Sum”)** and the only one among these top-5 targets which presents no opposing rules (**Supplementary Table 4, - “Targets selection”, column:”Contradictory_Sum”**). Finally, it has been found to interact with PTEN^52^ which is part of the PI3K/Akt pathway that plays a major role in cancer progression and has presented significant opportunities for the implementation of treatment strategies^53^.

Consecutively, POF1B was selected as the best target immediately after MAGI3 regarding strategy rules (*ii*) and (*iii*) (**Supplementary Table 4 - “Targets selection”, column:” Confirmatory_Sum”)**. POF1B’s expression was reported in malignant tumours regardless of their origin^54^ although its role in cancer is not well understood. Furthermore, POF1B is involved in the regulation of actin cytoskeleton,^55^ which has been shown to regulate the intracellular accumulation of doxorubicin and modify the resistance against the drug in osteosarcoma cells^56^.

Finally, PDIA3 although further down the list was selected because it has been shown to positively regulate the mTORC1 complex assembly and signalling^57^ and its inhibition has also been shown to increase sensitivity to ionizing radiation and chemotherapeutics^58^.

The selected genes were silenced in the A549 (lung carcinoma), NCI-H1299 (lung carcinoma derived from metastatic site), MCF7 (breast adenocarcinoma derived from metastatic site) and Saos-2 (osteosarcoma) cell lines treated with Doxorubicin (**Methods - Experimental rule verification**). A similar set of experiments was performed using Taxol as MAGI3 over-expression was found to be strongly associated with Taxol resistance (**Methods - Experimental rule verification**). Drug sensitivity was estimated as IC_50_ values in MTT assays before and after these treatments, while cell viability was not influenced by the genetic silencing of the examined genes (**Supplementary Figure 2**). Notably, in all cases siRNA treatments led to a significant sensitization of the examined cells to doxorubicin as predicted, with MAGI3 down-regulation showing a similar performance also in taxol sensitization experiments (**Figure 4e**, **Supplementary Table 4**). Analysis of TCGA gene expression data provides further insights on the potential use of the aforementioned genes in clinical practice for cancer types that frequently over-express the above genes (**Figure 4d**, **Supplementary Table 3**). Specifically, chromophobe renal cell carcinoma and breast invasive carcinoma present elevated MAGI3 mRNA levels in 53% and 50% of cases respectively. POF1B mRNA levels were found to be elevated in 99%, 97%, 82%, 76%, and 63% of rectum adenocarcinoma, colon adenocarcinoma, stomach adenocarcinoma, stomach esophageal carcinoma and esophageal carcinoma respectively. Finally *PDIA3* mRNA levels were found to be elevated in 62% and 54% of prostate adenocarcinoma and uterine corpus endometrial carcinoma respectively.

**Figure 4:**
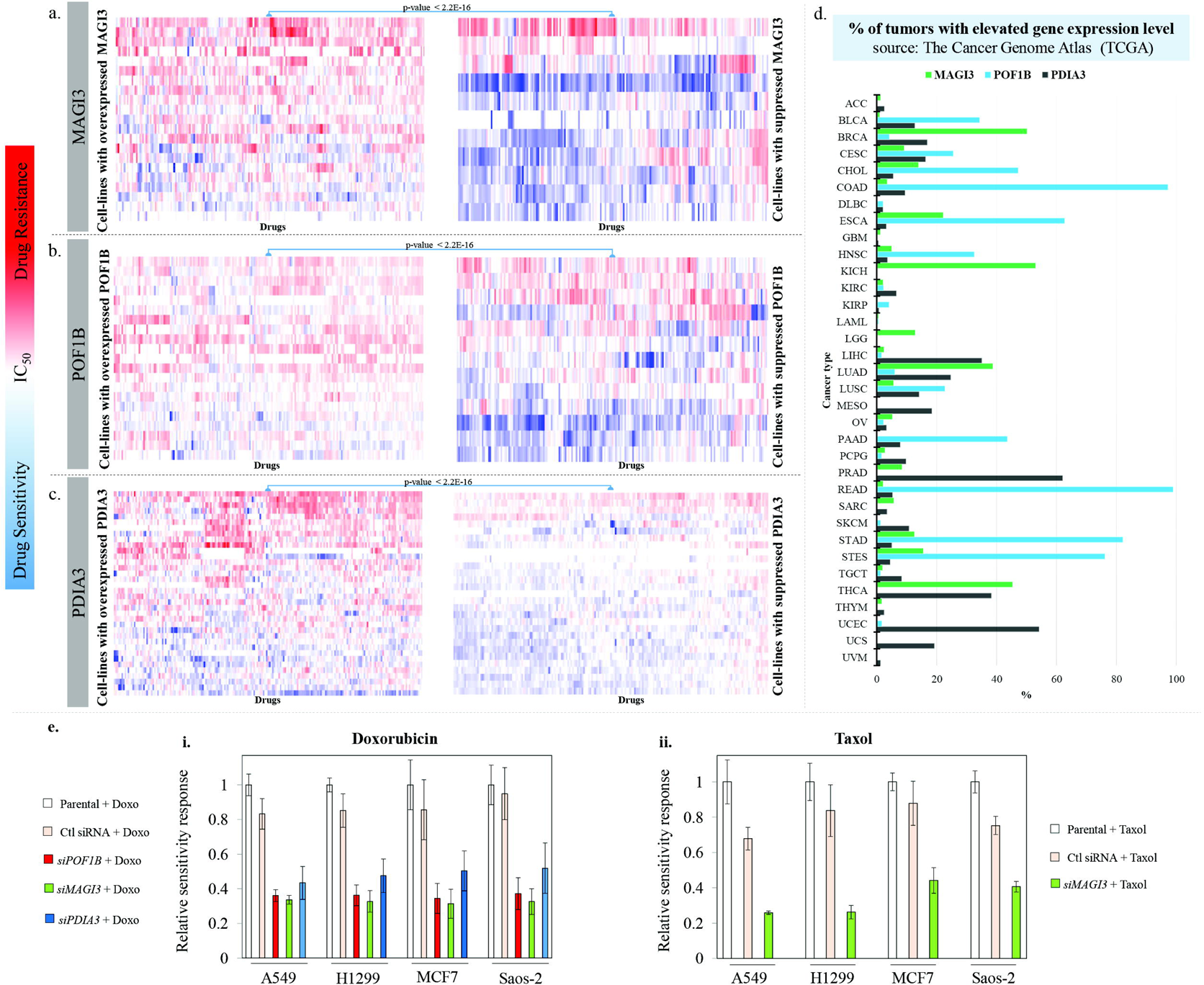
Experimental Validation of novel si-targets identified by the pipeline. **a)** IC_50_ heatmaps of drugs for cell-lines that over and under-express MAGI3. **b)** IC_50_ heatmaps of drugs for cell-lines that over and under-express POF1B. **c)** IC_50_ heatmaps of drugs for cell-lines that over and under-express PDIA3. **d)** Percentage of TCGA cases per tumour type which express MAGI3, POF1B and PDIA3 one standard deviation above and below the mean. **e)** Relative IC_50_ levels (measured as µg/ml), as estimated by the MTT-assay **(Supplementary Table 4)**, of the cell lines A549, H1299, MCF7 and Saos-2 when treated with *i)* Doxorubicin in combination with silencing of MAGI3, POF1B and PDIA3 compared to the IC_50_ levels of the cells when treated with the drug alone and *ii)* Taxol in combination with silencing of MAGI3 compared to the IC_50_ levels of the cells when treated with the drug alone. All IC_50_ levels were visualised relative to the IC_50_ levels of the cells treated only with Doxorubicin or Taxol respectively. P-values were derived by the non-parametric two-tailed Wilcoxon test.

### 4. Prediction of Drug-Response

*4a. Train & Test datasets for Machine Learning.* To predict drug response through machine learning, we split the main data-set into two subsets, referred to as training set and test set and consisting of approximately 2/3 and 1/3 of the main data-set, respectively. The detailed description of the sets construction is presented in the relevant **Methods section - Prediction of drug-response (Supplementary Figure 3**).

*4b. Machine Learning.* For our drug response classification framework, we applied DLNN^11^ enhanced by Bagging Ensemble Learning^59^. Although its performance has not been tested in drug response prediction, we selected the Deep Learning Framework because it has redefined the state-of-the-art in many applications ranging from image recognition to genomics^11^. In particular we chose to use the open-source DLNN framework provided by H2O.ai (http://www.h2o.ai/). The H2O.ai is a cluster ready framework, which allows for the machine learning part of our pipeline to be readily deployable to a high performance-computing environment. In order for machine learning to be able to perform well on the blind-set (test-set), it is critical to select only the most relevant features for training the classifier, for example features that are highly correlated with drug response. If a large number of irrelevant features are used for training, the classifier will be trained on noise, and although it will produce excellent results on the training set, it will perform poorly on the blind-set. This problem is referred to as over-fitting and in the application of “omics” information, where the number of features (in our case, gene expression, mutation status, etc.) vastly outnumbers the total number of cases, over-fitting is inevitable. To further address this problem for DLNNs we only used activation functions that utilize dropout^60^ which has been shown to effectively prevent neural networks from overfitting.

Previous reports have used elastic-net (as discussed above)^61^ as a feature reduction technique. To address this challenge and select the most relevant features, we utilised the rule-set generated by the Association Rule Mining, as performed on the training-set alone (**Supplementary Table 5 – Training-Set Rules**). The DLNN classifiers (one classifier per drug and per drug-response-state) were trained on the training set constructed from features selected by the Association Rule Mining procedure and respective drug-responses (**Supplementary Table 5 - Classification_training_features**). Each classifier’s performance was then assessed on a blind-set (the test-set), where we provided only the specific features upon which the classifiers predicted the drug response, which was then compared to the actual drug-response value. In order to compare the performance of the DLNN over other well-established machine learning methods, we repeated the classification task utilising RFs, BMMKL and shallow Neural Networks (NN). Both non-linear approaches, RF and BMMKL are the state-of-the-art classification frameworks providing top prediction performance in a similar drug response prediction setup^17^. RF is a highly adaptive tree-based machine learning tool, that has been applied for prediction and classification for genomic data, and unsupervised learning^62^. BMMKL utilizes kernelized regression to represent the relationships between features and Bayesian inference for learning the model^63^. Finally, all the aforementioned machine learning frameworks were trained over scrambled training and test sets (decoys) as negative controls hence as a means to quantify the effect of noise on classification (**Figure 5, Supplementary Table 5**). Technical details are presented in the respective Methods Section.

**Figure 5:**
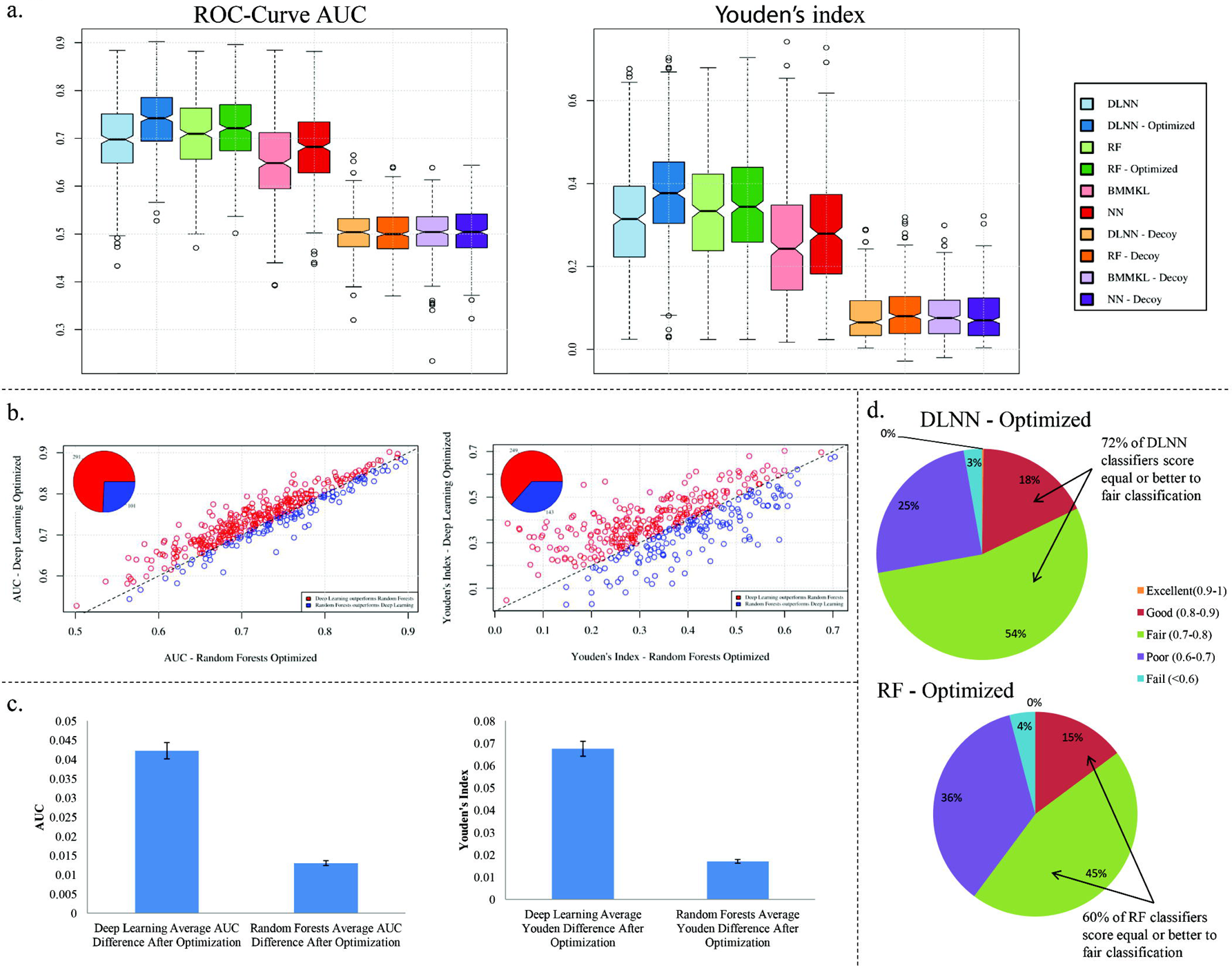
Comparison of prediction performance between DLNN, RF, BMMKL and NN. **a)** Classification performance assessment according to ROC-Curve Areas Under the Curve (AUC) and Youden’s Index for the machine learning frameworks. **b)** AUC and Youden’s Index scatterplots for optimized DLNNs and Random Forests per drug. Pie-charts within the scatterplots indicate the actual numbers of *i)* DLNN classifiers that outperform the respective Random Forests classifiers (red) and *ii)*Random Forests that classifiers that outperform the respective DLNN classifiers (blue). **c)** Bar plots with standard error bars of the improvement of classification performance after optimization for DLNNs and Random Forests. **d)** Percentage of classifiers that have scored excellent, good, fair, poor and random classification performance based on the ROC-Curve AUC for optimized DLNNs and Random Forests. All performance metrics were measured on the test set.

By using the genes involved in the association-rules as features and the DLNN as a machine-learning framework, we constructed classifiers capable to predict whether a cell-line would be sensitive or resistant to a given drug, based on its molecular profile. In agreement with a previous study^5^, since this dataset is comprised of many cancer cell-lines from different tissues of origin, we observed that the vast majority of predictive features are gene expression levels. Additionally, we noted that the information of tissue of origin significantly improved the prediction performance. Our pipeline produced a total of 392 classifiers corresponding to 196 drugs, each with two responses (sensitivity and resistance) for each one of the four classification frameworks, namely DLNNs, RFs, NNs, BMMKL plus all the aforementioned frameworks running over scrambled training and test sets (**Methods - Prediction of drug-response**). The DLNNs were initially used without any model optimization while Random Forests underwent a certain degree of model optimization for each case. Finally, suggested settings, according to the relevant literature, were used for NNs and BMMKL (**Methods - Prediction of drug-response**). We did not produce classifiers for all the drugs because we excluded the ones than had more than 20% of their response values missing. To evaluate the classification efficiency of our classifiers, we applied a series of metrics, namely Area Under the Curve (AUC) of the Receiver Operating Characterstic (ROC)-curve, Youden Index, Sensitivity, Specificity, Accuracy, Positive Predictive Value (PPV), Negative Predictive Value (NPV) and False Positive Rate (FPR). The results obtained for all tested classifiers are reported in **Supplementary Table 5.** The classification performance of the tested classifiers was based upon the ROC-curve AUC^64^ & Youden’s Index^65^ (**Figure 5**) for the following reason. The ROC-curve AUC is a metric that integrates true and false positive classification rates and does not depend on the selection of a discriminating threshold, while the Youden’s index is an established index for rating diagnostic tests combining sensitivity and specificity metrics. Youden’s Index along with all the remaining performance metrics, apart from the AUC, depends on the selection of a discriminating threshold. We therefore decided to use the threshold that maximised the Matthews correlation coefficient,^66^ which is a performance metric unbiased towards unbalanced data-sets such as our own. The performance of all classifiers can be seen in **Figure 5a** and is recorded in detail in **Supplementary Table 5.** Although Bayesian multitask learning was the top performer in Costello et al^17^, in our study it achieved the lowest performance. When we examined the details of the aforementioned work we noticed that the Bayesian multitask learning made extensive use of prior knowledge built in the training process in the form of additional features, while the random forests model used only the available data. The utilization of prior knowledge seems to be the reason for the achievement of top-performance Bayesian multitask learning. This fact is confirmed by a recent study which demonstrated that utilization of prior knowledge contributes to the improvement of the prediction process^67^.

In a preliminary comparison Random Forests slightly outperformed the initially un-optimised DLNNs. We hypothesized that this is due to the fact that the Random Forests framework is built to work efficiently with minor degree or even no model optimization^68^, while DLNNs are extremely complex models and model architecture is essential for the framework to achieve its full potential^10, 11^. To test this assumption we performed model optimization for both Random-Forests and DLNNs and compared the outcomes (**Figure 5**, **Supplementary Table 5**). More specifically, for each drug/state and for each of the two machine learning frameworks we performed grid search using the same training and test sets as in the prior analysis and each time we kept the model that achieved the best ROC-Curve AUC on the test set. We did that in order to monitor whether there would be a considerable improvement in the classification performance of the DLNNs and minor improvement in the classification performance of Random Forests as initially hypothesised. It must be mentioned that, for Random Forests, we performed a model optimisation procedure (**Methods - Prediction of drug-response**) that according to the framework’s internal mechanics maximises the possibilities for an optimum or near-optimum model. On the other hand the parameters of a DLNN model are numerous requiring for thousands of models to cover all possible combinations. We therefore selected a small set of parameters for implementing only a 3-layered DLNN with three different dropout-based activation methods to avoid over-fitting (**Methods - Prediction of drug-response**) and we performed a discrete random sampling of only 150 models from the available model sample space each time due to computational limitations given that DLNN models are extremely expensive to train in terms of computational resources and CPU time. Therefore, in the case of DLNNs model optimization was by no means as thorough as in Random Forests, meaning that the chances of obtaining a near-optimum model were significantly worse in comparison to Random Forests. The results confirmed our initial hypothesis (**Figure 5c**, **Supplementary Table 5**) as the average improvement of classification performance in terms of ROC-Curve AUC was 4.2% and 1.3% for DLNNs and Random Forests respectively and in terms of the Youden’s index 6.7% and 1.7% respectively. After optimisation the DLNNs clearly deliver top classification performance with both performance metrics (*p*-value < 0.05) (AUC & Youden’s Index) (**Figure 5 a,b**, **Supplementary Table 5**). According to the widely accepted AUC-based classification quality grading scale, classifiers that produce AUCs 0.90 - 1 are considered excellent, 0.80 - 0.90 are good, 0.70 - 0.80 are fair, 0.60 - 0.70 are poor classifiers, while classifiers with an AUCs below 0.6 are considered failed or random classifiers^64^. Our pipeline produced 392 classifiers corresponding to 196 drugs, each with two responses (sensitivity and resistance) (**Methods - Prediction of drug-response**). Out of these trained and tested optimized DLNN classifiers, only one was excellent, 18% good, 54% fair, 25% poor and 3% random classifiers as opposed to 0%, 15%, 45%, 36% and 4% for the optimized Random Forests using the AUC classification quality grading scale (**Figure 5d**, **Supplementary Table 5).** More specifically 60% of the Random Forest classifiers scored equal to or better than a fair classification (AUC>0.7) whereas 72% of the DLNN classifiers achieved the same performance, indicating a superior performance of DLNN over Random Forest classification quality.

### 5. Drug-Clustering

#### 5a Drug clustering based on Jaccard distances

Combining drugs against multiple targets belonging to inter-linked or overlapping signalling cascades are strong candidates for presenting synergistic effects^69^. Our aim was to create a clustering scheme based solely on the presence of specific genes derived from the rule-set connected to a specific drug-response. To this end, we produced two individual clustering schemes, one for drug-sensitivity (**Figure 6**) and one for drug resistance (**Supplementary Figure 4**). Subsequently, we focused on the analysis and presentation of the most stable clusters highlighted in red without this meaning that the remaining clusters are to be rejected. All dendrograms can be accessed in HTML format in the folder “/Figures/Dendrograms/” at the GitHub repository **(Methods - Data Availability**). For details on clustering and determination of stable clusters refer to **Methods - Drug-clustering**.

**Figure 6:**
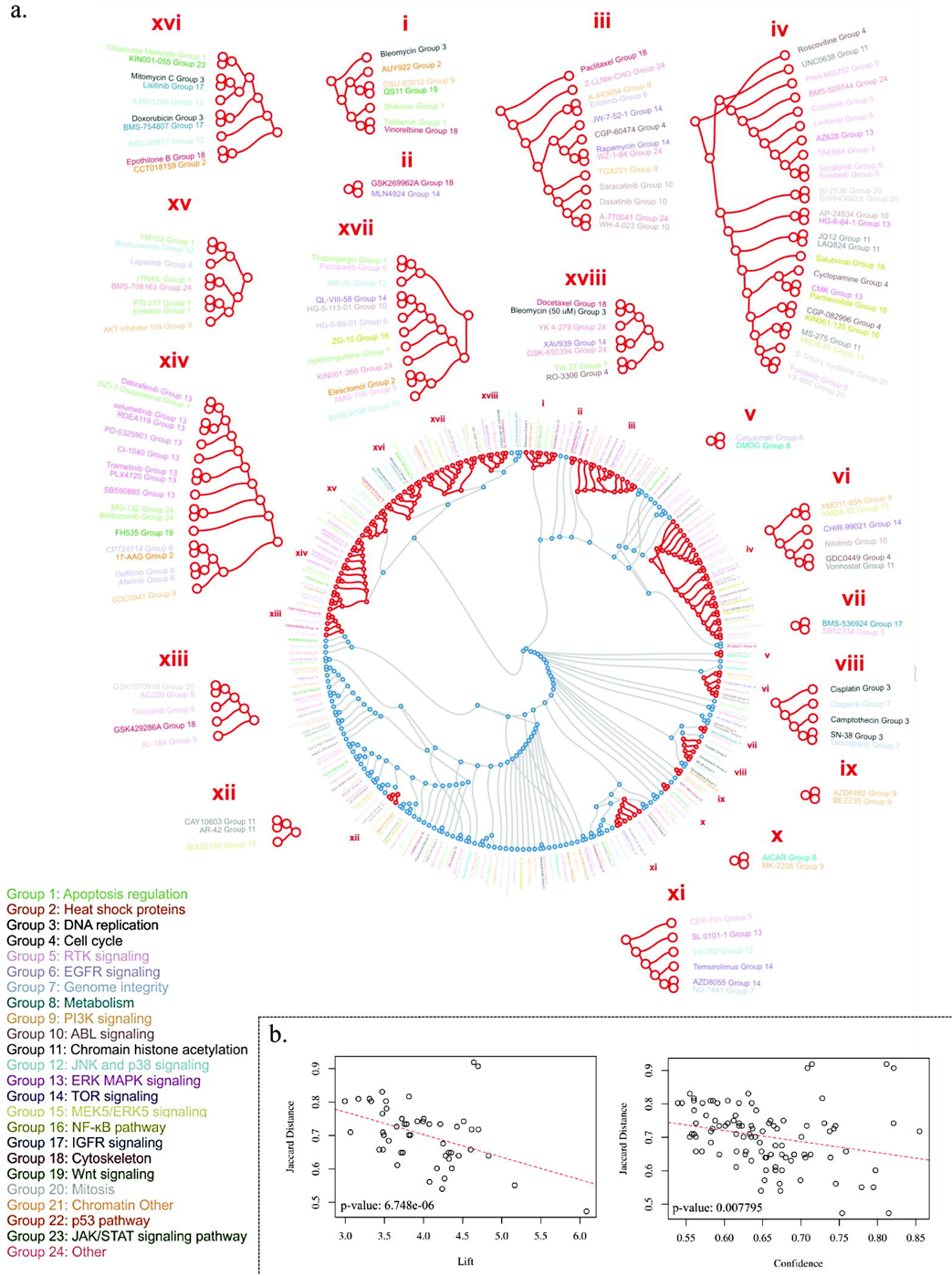
Drug sensitivity clustering. **a)** Drug sensitivity clustering based on the similarity of genes involved in the association rules with the sensitivity state of each drug. **b)** Correlation of Jaccard-distance with rule confidence and lift for drug-to-drug association rules (sensitivity state) (**Supplementary Table 8 - “Sens_Sens”**). Drugs are colour coded by drug type. Detailed information regarding the groups and the associated drug targets is located in **Supplementary Table 2**. Red clusters represent stable clusters.

By examining the two circular dendrograms, it was clear that they bore no-resemblance to each other meaning that the differentiated genes in the cell-lines that are resistant to a specific drug are dissimilar from the differentiated genes in the cell-lines that are sensitive to the same drug. To confirm this observation, we utilised the features used for classification training (**Supplementary Table 5 – “Classification_training_features”**), calculated the overlap between all possible drug-state combinations, (**Supplementary Table 6 - “Genes overlap”)** and determined whether the magnitude of the overlap was random or over/under represented (**Supplementary Table 6 - “p-values”, Supplementary figure 5 and 6**). Given the size of the sensitivity and resistance gene-sets for each drug, this calculation was accomplished by Monte-Carlo simulation, as described in the **Methods - Drug-clustering**. This calculation highly correlated with the produced dendrograms, as drug-response-states, that cluster together, presenting statistically significant over-represented gene-set. We isolated all of the above information related to the sensitivity and resistance states only for each drug as presented in **Supplementary Table 6 - “comparison sens & res per drug”.** We observed a very low overlap among the gene-sets involved in the sensitivity and resistance states of any given drug (**Supplementary figure 5 and 6**), indicating that the pathways involved in sensitivity and resistance for any given drug are diverse, which is in perfect agreement with our prediction strategy of using different models for predicting sensitivity and resistance for each drug. Examining closely the sensitivity dendrogram clustering structure, we noted that it was highly relevant to the drugs target (**Figure 6**), indicating that drugs with the same target tended to cluster in close proximity to one another. This observation was anticipated as the gene product targeted by a drug is part of the sensitivity predictive gene-signature of that particular drug (**Figure 3, section 3a – Proof of concept 2).** Additionally, the sensitivity-status clustering appeared to capture the broader relationships among the drugs. For instance, we observed that there was a branch populated not only by BRAF, but also by MEK inhibitors, which practically belong in the same signalling network (**Figure 6 – Cluster XIV**)^70^. In contrast, when we examined the resistance dendrogram (**Supplementary Figure 4)**, we observed that clustering is less relevant to the drug targets in comparison to the sensitivity dendrogram meaning that the molecular cascades implicated in drug resistance are diverse from the ones being targeted by the drug. To determine whether drug clustering translates to highly correlated activity of closely clustered drugs (sensitivity dendrogram) across the cell-lines, we extracted the drug-to-drug rules from our total rule-set (**Supplementary Table 8 - “Sens_ Sens**”) to examine whether sensitivity responses of the cell lines to certain drugs are correlated to others. We clearly observed that the rules with the largest support had their Lift and Confidence values inversely correlated to the clustering distance in the dendrogram. This means that drugs which are connected in these rules tend to cluster closer in the sensitivity dendrogram **(Figure 6b),** implying that the corresponding genes are involved mechanistically in producing a drug sensitive environment.

#### 5b. Suggestion of a Rule for the determination of drug partners with high potency

Based on drug clustering, we propose a drug-pair selection strategy for combined therapeutic approaches using the following rule: Candidates with a high probability for presenting synergistic effect are those that: a) target different molecules, b) are located close together in the sensitivity dendrogram with their proximity also confirmed in the drug-to-drug association rules, and c) cluster as far-away from each other in the gene-based resistance dendrogram as possible.

The last rule helps us select drugs with a synergistic potential due to the high overlap that the corresponding, targeted by the drugs, signalling cascades demonstrate^71^. This is clearly depicted by their proximity in the sensitivity dendrogram. Concurrently for an effective treatment we need **the same drugs** to present highly diverse resistance-related pathways, as indicated by their larger distance at the resistance dendrogram. The latter renders the cancer cell harder to acquire the required molecular alterations for dual drug resistance.

#### 5c. Proof of concepts supporting the Drug-Partner Rule

##### Proof of concept 1

In Clusters III, VI and XVII of the drug synergy dendrogram the ABL and TOR inhibitors are closely situated **(Figure 6),** implying a synergistic effect. Indeed, increased synthetic lethality has been reported in leukemias by administrating together ribavirin (mTOR inhibitor) and imatinib (ABL inhibitor)^72^. Along the same lines, combining mTOR inhibitors with PI3K inhibitors (derived from Clusters III and VI) had a greater anti-leukemic effect than monotherapy^73^.

##### Proof of concept 2

Drugs in cluster VIII, involved in modulation of DNA replication and genome integrity maintenance, have been shown to be more effective when administrated concurrently. For instance, combination of olaparib [PARP (Poly-ADP Polymerase)-inhibitor blocking DNA repair - genome integrity] with camptothecin (Topoisomerase-I inhibitor – blocking DNA replication) renders cancer cells radiosensitive^74^. Likewise, combining olaparib with cisplatin (inhibitor of DNA replication) or SN-38 (Topoisomerase-I inhibitor), has been shown to be efficient in treating lung^75^ and colon^76^ cancer, respectively

##### Proof of concept 3

Therapeutic agents in cluster XIV, targeting the ERK, MAPK and EGFR signalling routes, display an additive effect when concurrently administrated^77, 78^. Examples include, colon cancer [Gefitinib (EGFR inhibitor) + Vemurafenib or Dabrafenib (BRAF inhibitors)]^79^ and melanoma [BRAF + MEK inhibitors – Group 13 of cluster XIV]^24^, as discussed earlier (**section 3a, Proof-of-concept 2**). Moreover, specific drugs found in this cluster, such as GDC-0941 (PI3K inhibitor) and PD-0325901 (MEK inhibitors) inhibit the growth of colorectal cancer cell lines more efficiently when added together^80^

##### Proof of concept 4

Among the drugs in cluster IV we observed grouping of HDAC (histone deacetylase) inhibitors with NF-κB inhibitors. Several studies reported that HDAC inhibitors act as modulators of NF-κB signalling^81^-^83^, suggesting that dual inhibition could appear promising in inflammatory driven cancer^84^. Similarly, in clusters IV and VI we observed a potential synergy between histone deacetylase inhibitors (HDACIs) and therapeutic agents affecting cell cycle signalling and development, such as the Hedgehog (Hh) pathway inhibitor cyclopamine or GDC- 0449. Increased HDAC6 expression was found crucial for optimal activation of Hh pathway and its inhibition affected the survival of medulloblastoma cells^85^. Moreover, dual inhibition of histone deacetylases and Hh pathway was shown to overcome resistance to single Hh targeting agent**^86^.**

Finally, we noticed that cluster IV is enriched with inhibitors targeting receptor tyrosine kinase (RTK) signalling. Among them we noticed AZ628, a BRAF inhibitor. Interestingly, RTKs are utilised by cancer as a means to resist BRAF inhibitors; therefore synergistic therapy with anti-RTK and anti-BRAF agents appears as an appealing therapeutic strategy to prevent BRAFi (BRAF-inhibitory) resistance^87^.

## DISCUSSION

We present an *in silico* pipeline that utilises a large cancer cell-line dataset (1001 cell lines) with diverse genomic features (>60,000 features) and responses to a diverse number of drugs (251), to extract knowledge in the form of easily interpretable rules and then, by combining these rules with the sate-of-the-art DLNN framework, to accurately predict drug responses. We also demonstrate that prediction of sensitivity and resistance responses must by handled by different models since the genes, hence the underlying molecular cascades that drive these responses are diverse. Lastly, we suggest a strategy, based on the drug sensitivity and resistance clustering to select the most potent candidates for drug-combination therapy.

Association rule mining is an efficient big-data ready algorithm that produces a framework of easily interpretable information in the form of simple rules that offer novel insights on the data, which in turn lead to generation of novel hypotheses. Although it could be argued that association rule mining is no different than univariate statistics, there are several points that justify our preference over this framework. Univariate statistics utilize hypothesis testing (t-test, Wilcoxon-test, chi-squared test etc.) to investigate, for instance, whether there is a real difference in the mean value between two groups or whether a group is enriched in one condition versus another, and the p-value is used as a measure of rejecting or accepting the null hypothesis. On the other hand, association rule mining creates all possible associations between variables and evaluates these association patterns with more than 30 different significance measures^19^ such as Support, Confidence, Lift, Gini Index, Entropy and others, each one providing different views on the data, hence greater flexibility to revealing hidden association patterns from complex datasets. Additionally, association rule mining provides the possibility for exploring deeper relationships among the variables, such as multi-way interactions to be detected and evaluated for significance; further enhancing the identification of hidden patterns from the aforementioned complex datasets.

Examples of rules validation and utilization such as the identification of ID1 as a potential biomarker for identifying patients that will benefit from treatment with PI3K inhibitors, are presented in a “proof of concept” manner representing a small fraction of possible interpretations.

Furthermore, the association rule framework was utilised to device an algorithm for identifying novel targets which through gene silencing or chemical inhibition would enhance the effectiveness of already established drugs. The algorithm was validated with complete success examining experimentally three newly identified targets for their sensitivity towards doxorubicin and taxol **(Figure 4).** This is a major step in the direction of personalized medicine scheme since it enables us to formulate custom made nano-particles, as previously described^88^, loaded with the appropriate drug and si-RNA molecules, based on the patients gene-expression profile, achieving maximum therapeutic efficiency.

Association rules were further exploited as a feature selection tool for predicting drug-response through machine learning. To the best of our knowledge, the current study is the first to demonstrate the effective utilization of the Deep Learning framework to predict drug response from molecular profiling data clearly outperforming, even when unoptimised, its predecessor, the Neural Networks. Although Hornik et al. proved that any complex function can be approximated by a single layer neural network^89^, in practice their approximation becomes inefficient as the dimensionality of the feature space increases and that is the reason why neural networks were forsaken by late 90s^11^. On the contrary, DLNNs which are multi-layered neural networks overcome this inefficiency and learn complex representations of multi-dimensional data with multiple levels of abstraction^11^. Furthermore, we demonstrated that after model optimisation, DLNNs clearly outperformed RF and BMMKL which are currently considered the state-of-art in drug-response prediction^17^.

Finally, we suggest a strategy, based on the drug sensitivity and resistance clustering to select the most potent candidates for drug-combination therapy. We provide a number of examples were drugs that cluster together in the sensitivity dendrogram act synergistically (**see results section 5**). Moreover, novel relationships between drugs are unrevealed that need further investigation for potential synergy (**Supplementary Table 7)**. For instance, one of the strongest relationships in the sensitivity cluster is between the drugs XMD14-99 [EPHB3 (Ephrin-B3) inhibitor] and JW-7-24-1 [LCK (Lymphocyte-specific protein tyrosine kinase) inhibitor] (**Figure 6, Supplementary Table 7 - “Jaccard_Sensitive”).** Although their synergy has not been tested in the lab, Jiang G et al., reported that in human leukemia cells, ephrin-B-induced invasive activity is supported by LCK^90^, implying that a combination scheme against them could be therapeutically effective. Another potent relationship was observed between Epothilone B (microtubule associated inhibitor) and CCT018159 [HSP90 (Heat Shock Protein-90) inhibitor] (**Figure 6**, **Supplementary Table 7 - “Jaccard_Sensitive”).**

Inhibition of this pair has been considered for Alzheimer’s treatment as synergy between tau (a microtubule-associated protein) aggregation inhibitors and tau chaperone modulators inhibitors (HSP90) and were shown to be effective^91^. On the topic of cancer treatment, Zhang et al., reported the anti-tumour activity of CDBT (2-(2-Chlorophenylimino)-5-(4-dimethylamino-benzylidene) thiazolidin-4-one), which is a dual microtubule and HSP90 inhibitor in a multi-drug resistant non-small-cell lung cancer model^92^. When considering which synergy pair to select for experimental validation we should also bear in mind the status of the specific pair in the Resistance Table (**Supplementary Table 7 - “Jaccard_Resistant”)** and de-prioritize it if the resistance relationship of the pair is strong. The ground for this criterion is the high overlap of genes involved in resistance development of the two drugs, which practically means that when the cancer cell acquires resistance to the first drug, will most probably acquire resistance to the second as well. As a final note, clustering was performed using the pan-cancer dataset, therefore tissue specific information regarding potential synergies is not available in the current version. We selected to analyze the pan-cancer dataset because the current size of the available dataset was a limiting factor for producing tissue specific synergistic trees, which of course is the next step for increasing the accuracy of the described strategy.

Our vision for the current work is the development of an expert decision support system where the molecular profile of the patient’s tumour would be uploaded and the system would predict the drug response for a large screen of drugs allowing clinicians, through an augmented medicine scheme, to select the best candidates for mono or combination therapy. Additionally, the effectiveness of these candidates could be further enhanced by silencing the proper genes as also indicated by the system. These therapeutic schemes would then be tested on patient-derived primary 3D cancer cell cultures^93^ and/or on xenograft models^94^. The most efficient combination would then be applied in the form of a therapeutic scheme directly on the patient with constant monitoring for administration of personalised dosing.

The power of the presented pipeline lies on the efficiency, expandability and ability to create easily interpretable rules of the Association Rule mining algorithm, and to the capability of Deep Learning to capture the complex heterogeneity of tumours. It can be further expanded by increasing the number of cancer cell-lines, including patient-derived cell-lines, as well as by increasing the number of therapeutic agents analysed by the system. Additionally, the system allows integration of other layers of “omics” information, including meta-genomics, proteomics, phospho-proteomics, interactomics and metabolomics, which will further enhance the prediction and drug-clustering schemes. We propose that the bioinformatic pipeline described is expandable and effective utilising state-of-the-art algorithms such as Association Rule Mining and Deep Learning and can effectively be applied in the rapidly developing “omics” era for devising personalised medicine schemes, as well as for biomarker and drug discovery.

## METHODS

All scripting, data-processing, statistical calculations have been performed with R-language for statistical computing^95^.

### 1. Datasets

The dataset compilation from “Genomics of Drug Sensitivity in Cancer” (GDSC - release 6) and “COSMIC Cell Line Project” (CCLP) was created by the R script “**script_make_data.R**”.

Tissue of origin and drug response data were obtained from: ftp://ftp.sanger.ac.uk/pub4/cancerrxgene/releases/release-5.0/gdsc_manova_input_w5.csv.

Gene mutation data was obtained from “**CosmicCLP_MutantExport.tsv**”, gene expression data was obtained from “**CCLP_CompleteGeneExpression.tsv**” and copy number variation data was obtained from “**CosmicCLP_CompleteCNA.tsv**”. All the aforementioned files were downloaded from http://cancer.sanger.ac.uk/cell_lines/download. More specifically, with respect to the molecular profiling data we included mutational status for 19384 genes, copy-number-variation status for the exons of 24858 genes and gene-expression status for 16445 genes. The gene mutation status is a factor consisting of 1 level, namely “Mut” that corresponds to all single point mutations apart from the silent ones. The copy-number-variation status is a factor consisting of two levels, namely “Gain” and “Loss” for gains and losses, respectively, while the gene-expression status is a factor that also consists of two levels (“over” and “under”) that correspond to z-scored gene expression levels greater and lower than two standard deviations from the mean, respectively. Finally the drug status is a factor consisting of two levels, namely “Resistant” and “Sensitive” that correspond to z-scored IC-50 levels greater and lower than one standard deviation from the mean, respectively. The R-Data object containing the matrix is stored in the file **MASTER_MATRIX.RData**.

### 2. Association Rule Mining

#### 2.1 Apriori Algorithm

To provide insights regarding the way the algorithm works we provide an example. Gene expression of Gene-A in our dataset has two levels, “over” and “under”. The **Apriori algorithm** will generate two features out of Gene-A gene expression, namely Gene-A=over & Gene- A=under. The rules come in the form of A => B. The feature A is considered to be the Left Hand Side (**LHS**) of the rule while the feature B the Right Hand Side (**RHS**). For the scope of the current study, we only kept the rules containing drug sensitivity features on the **RHS.** The algorithm can also be utilized to mine for more complex association rules containing interactions on the **LHS** in the form of A, B => C which is a two-way interaction, being able to go as deep as the data-set and the computational resources permit.

There are three basic metrics utilized by the algorithm in order to describe the power and significance of the rules. These metrics are **Support**, **Confidence** and **Lift**. **Support** is the frequency of the rule occurrence in the total dataset. **Confidence** is the frequency of rule occurrence in the cases of the dataset fulfilling the **LHS** of the rule. Finally **Lift** is a measure of significance. For the simple rule 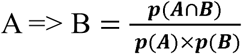, which, based on probability theory will be equal to 1 if the features A & B are independent. For dependent features the value of **Lift** will be greater than 1 and the value being proportional to the power of the association. In order to run the Apriori algorithm, the user has to define minimum support and confidence values below which all rules are discarded, plus the number of allowed interactions in the **LHS**. We initially ran the algorithm by setting a minimum support and confidence of 0.58%, corresponding to just 6 out of the total of 1001 cell-lines allowing for no interactions (1-way: A =>B), which is the minimum our computational resources permitted. Finally, we ran the Apriori algorithm at minimum support and confidence levels of 1.02% (due to limitations in computational resources) allowing for one interaction (2-way: A + B => C).

#### 2.2 Dynamic Thresholding

The Apriori algorithm was run on a permuted version of our initial dataset (**MASTER_MATRIX_PERMUTED.RData**), which was produced by randomly shuffling each individual column of the dataset. The permutated matrix was produced with the script **script_make_data.R**. We initially ran the algorithm with the aforementioned support and confidence values on the permuted dataset and we determined the Lift threshold that would control the false discovery rate at less than 5%. We noted, however, that for each different set of support and confidence values belonging to our actual rules, there was a different lift threshold for FDR<5% if the Apriori algorithm had run on the permuted dataset with that set of support and confidence values as the minimum support and confidence parameters of the algorithm respectively. We therefore adjusted our thresholding determination with a method we call **Dynamic Thresholding**. Specifically, for every unique set of support and confidence values, we ran the Apriori algorithm on the permuted dataset using these values as the minimum support and confidence required by the algorithm, and we then determine the Lift threshold for which FDR=5%. After the completion of that process we evaluated each one of our actual rules based on its Lift value; if above of the specific threshold, the rule was accepted as significant, otherwise it was rejected. Both 1-way and 2-way rules where filtered keeping only the significant rules (FDR<5%). The significant rules are available in **Supplementary Table 1**. The rule-set constitutes a novel meta-data-set, which can be utilized for knowledge extraction as per the proof s of concept that follow in the current text.

The implementation of the Apriori and Dynamic Thresholding algorithms can be found in the script “**script_dynamic_thresholding.R**”.

#### 2.3 Group-wise rule visualization

The group wise Association Rules visualization presented in the current study utilizes k-means clustering in order to visualize data with high dimensionality and high scarcity and are described in detail in Hahsler et al. (2011)^96^ and is implemented in script “**script_rules_visualize.R**”.

### 3. Experimental rule verification through gene silencing

#### 3.1 Cell lines

Cell lines were maintained in DMEM (Invitrogen) with 10% FCS (Invitrogen), 2mM l-glutamine (Invitrogen), and 100µg/ml penicillin and streptomycin (Invitrogen) at 37°C and 5% CO2.

#### 3.2 siRNA transfections

*POF1B* (#127487-9), *MAGI3* (#123257-9), *PDIA3* (#107677-9) and Silencer® Negative Control #1 (4404021) (Thermo Scientific) siRNA gene silencing was performed as described^2^, following also the manufacturer’s instructions.

#### 3.3 RNA isolation, cDNA preparation and real time (RT)-PCR

RNA was extracted with the RNeasy Mini Kit (#74104, Qiagen). cDNA generation and real-time reverse transcription PCR (RT-PCR) analysis was run as described before^2^. Primer sequences and annealing temperatures are provided in **Supplementary Table 4**. Results are presented as n-fold changes versus the values of the control-siRNA treated sample. Mean value was calculated from three independent measurements.

#### 3.4 MTT Assay

Cytotoxicity was estimated by the MTT (3-[4,5-dimethylthiazol-2-yl]-2,5-diphenyltetrazolium bromide) assay, as previously described^97^. Briefly, the cells were plated in 96-well, flat-bottomed microplates at a density of approximately 15,000 cells/cm2, in DMEM containing 10% FBS. Twenty four hours after the plating, the medium was changed with a new one containing the chemotherapeutic agents, while one hour earlier the corresponding siRNAs had been administered. After 72 hours of incubation, the medium was replaced with MTT dissolved at a final concentration of 1 mg/ml in serum-free, phenol-red-free DMEM, for a further 4-hour incubation. Then, the MTT formazan was solubilized in isopropanol, and the optical density was measured at a wavelength of 550 nm and a reference wavelength of 690 nm. A mean value was calculated from three independent experiments.

#### 3.5 Cell viability Assay

Cell viability was estimated using the CytoSelect™ Cell Viability and Cytotoxicity Assay (Cell Biolabs, Inc.; Cat No CBA-240) following the manufacturer’s instructions. Results were averages from three independent experiments.

### 4. Prediction of drug-response

#### 4.1 Training & Test Sets

The training and test sets were created by the R script “**script_make_data.R**”. The original z-scored gene expression levels were restored to the total matrix and the Training and Test subsets were constructed by performing blocked randomization on the original matrix. The blocking factor was the tissue type, and two thirds of the cell lines from each tissue type were randomly assigned to the Training sets and the remaining one third to the Test set. The ratios were always rounded in favour of the Training set. If there were only two cases for a particular tissue type then they were evenly split between the Training and Test sets and if there was only one case, it was assigned only to the Training set. Additionally, the gene-expression factors were replaced with the original z-transformed gene expression levels. The Training set consisted of 669 and the test set of 332 cell-lines (**TRAIN_GE_NUM.Rdata**, **TEST_GE_NUM.Rdata**, **Supplementary Figure 3**). In both sets there were several cell-lines lacking gene-expression information. These cell lines were removed.

#### 4.2 Feature Selection

The Apriori – Dynamic Thresholding algorithm, as described above, ran on the Training set alone at minimum support and confidence levels of 0.45%, in order to produce rules having no feedback from the Test set used for measuring the classification performance (**Supplementary Table 5 - Train-Set Rules**). For every drug and for each different drug response (Sensitive or Resistant) the genes present in the respective relevant rules having Support values greater than the support-values 1st quantile level were grouped and used as drug-state-specific feature subset along with the information on the tissue of origin (**Supplementary Table 5 - “Classification_training_feature”**) for training as many individual classifiers. The 1st quantile condition was used because it provided better predictions as measured from intra-training-set k-fold cross-validation utilizing the ROC-curve AUC as the performance metric. For the total of the 196 drugs which contained less than 20% missing values in their drug response variable, each one having two states, Sensitive and Resistant, 392 classifiers we constructed for each machine learning framework tested.

#### 4.3 Deep Learning

Deep Learning Neural Networks (DLNN) were constructed using the H2O.ai platform [http://www.h2o.ai/] each consisting of 3 hidden layers with 300 neurons in each layer using Maxout with Dropout as the activation function and class balancing. Further parameters for the DLNNs were: number of epochs=300, input dropout ratio=0.1, hidden dropout ratio=0.3. Internal performance metrics were acquired using 3-fold cross-validation. Deep Learning is implemented in the script “**h2o_DLNN.R**”. The H2O.ai platform was selected because it provides a cluster-ready framework for immediate and on-demand scaling-up.

Each DLNN was utilized in a 200-feature bagging-ensemble learning scheme. For each feature set, multiple training rounds, enough to over-sample the feature-set 3 times were performed. In each training round 200 features were randomly selected. If the total feature-set number was lower than 200, only 1 training round was performed with the full feature-set. At the end of each Training round, the DLNN was asked to predict the probabilities for the Test set. The Test-set predicted probabilities from each Training round were averaged to produce the final Test-set predicted probabilities (row-wise, hence for each Test-set cell-line) using a weighted averaging scheme, the weight being the ROC-curve area under the curve (AUC) calculated from the Training step of each round based on 3-fold cross-validation. After the completion of the Test-set prediction, the classification performance was measured by calculating the Area Under the Curve (AUC) of the ROC-curve, Youden’s Index, Sensitivity, Specificity, Accuracy (ACC), Positive and Negative Predictive Values (PPV & NPV) and False Positive Rate (FPR) of the prediction by utilizing the ROCR-package^98^ **(Supplementary Table 5).** For all the calculations of the aforementioned metrics apart from the AUC, the selected class-discriminating threshold was the one maximizing the Matthews correlation coefficient^66^ which is known to be the most appropriate measure for imbalanced classes.

#### 4.4 Random Forests

Random Forest classifiers (RF) were constructed again using the H2O.ai platform. Each classifier consisted by number of trees equal to half the number of features utilized for training^99^. Class balancing and 5-fold cross validation was used. The exact parameters can be found in “**h2o_RF.R**”.

#### 4.5 Bayesian Multitask Multiple Kernel Learning

Description of Bayesian Multitask Multiple Kernel Learning (BMMKL) method can be found in Costello et al.^17^. After Association Rule Mining was utilised for feature extraction as described above, classification views were computed in the form of Gaussian kernels (for numerical data) and Jaccard similarity matrices (for categorical data). Jaccard kernels for data having multiple categories i.e. CNV “Gain” and “Loss” were calculated separately for each state. For missing values, the minimum Jaccard similarity coefficient found the matrix was used. The following four kernels (views) were used as learning input: gene expression, mutation status, CNV_gain, CNV_loss. BMMKL algorithm was run with default initial parameters. The classification performance for each classifier was measured as described above. The **BMMKL_kernels_bagging.R** script performs data preparation (rules extraction, bagging, compilation of the test and training sets in the form of kernelized views), classification and 3-fold cross-validation of the model. The Bayesian classifier utilizes train and test functions, accessible through the Scripts “**BMMKL_supervised_classification_variational_train.R**” and “**BMMKL_supervised_classification_variational_test.R**”.

#### 4.6 Neural Networks

Shallow neural networks (NN) were constructed by the Deep Learning framework of the H2o.ai platform using only one hidden layer. The remaining parameters for optimal performance were acquired from Menden et al^100^. The script can be found at ‘**h2o_NN.R**’.

For RFs, BMMKL and NNs we utilized exactly the same bagging scheme as for the DLNNs. The classification performance was measured in all cases as described in the DLNN section **(Supplementary Table 5).**

#### 4.7 Classification on permutated data (Negative Control)

The rows of the training and test sets were shuffled independently for each column, producing the permutated training and test matrices (‘**TRAIN_GE_NUM_PERMUTED.RData**’, ‘**TEST_GE_NUM_PERMUTED.RData**’). Training and classification were then performed on the permutated training and test sets respectively for each machine learning framework.

#### 4.8 Model Optimisation

Model Optimization was performed without bagging due to extremely long run-times.

*Deep Learning:* Given that no specific guidelines exist in DLNN hyperparameter optimization we empirically chose the following parameters: (a) all three available activation functions with Dropout to minimize overfitting (MaxoutWithDropout, RectifierWithDropout and TanhWithDropout), (b) two input dropout ratios: (0.5, 0.1), (c) four sets of hidden dropout ratios: (0.1, 0.1, 0.1), (0.5, 0.5, 0.5), (0.1, 0.5, 0.1), (0.5, 0.1, 0.5), (d) seven sets of neuron numbers in each of the three hidden layers (only 3-hidden layer DLNNs were utilised): (100,100,100), (200,200,200), (300,300,300), (100,200,300), (300,200,100), (300,200,300), (200,300,200) and finally (e) two training epochs: 100 and 200. This grid search totals 336 models per drug-state and the search strategy was Random-Discrete for 150 models; hence only 150 models randomly selected of the total model features space was evaluated each time.

*Random Forests:* Based on common practice and empirical rules^62, 68^ various number of trees and number of features randomly sampled as candidates at each split were selected for the optimisation process. Specifically possible number of trees were (i) equal to the number of features, (ii) half the number of features, (iii) a quarter of the number of features (iv) 3 quarters of the number of features (v) 1000, 3000 and 5000 trees. Possible number of features randomly sampled as candidates at each split were (i) the square root of the number of features, (ii) half the square root of the number of features and (iii) twice the square root of the number of features. This grid search totals 21 models per drug-state and the search was exhaustive; hence all possible combinations were evaluated.

The script ‘**h2o_DLNN_RF_Optim.R**’ was used for the optimisation of both DLNN and RF.

### 5. Drug-clustering

#### 5.1 Drug-clustering based on Jaccard Distance

Clustering were based upon the genes involved in the sensitivity or resistance state of each drug as extracted from the 1-way rules of the aforementioned Apriori algorithm (**Supplementary Table 6**). The top 100 rules ranked by support and top 100 rules ranked for Lift for each drug and response, were combined. These rules were then converted into a binary matrix where 1 denotes the presence of a rule and 0 denotes the absence. From this matrix a dissimilarity matrix was calculated using the vegdist function and the Jaccard index^101^ from the R package “vegan”^102^. Hierarchical cluster analysis was then performed using the hclust function from the R package stats using the average clustering method^95^. Clusters were evaluated for stability with the pvclust R-package^103^. The resulting cluster denrodrograms are displayed in a circular format using D3: Data-Driven Documents^104^. The most stable clusters (p<0.05) are marked in red. The html versions of the two dendrograms can be accessed through GitHub “/Figures/Dendrograms”.

#### 5.2 Monte-Carlo Simulation

*Drug Resistance – Sensitivity gene-set overlap:* For every drug and each drug state (Sensitive or Resistant) the number of genes participating in statistically significant association rules was measured (**Supplementary Table 5 – “Classification_training_features”**). The gene-set overlap for all the combinations of the drug/drug-state pairs was also measured (**Supplementary Table 6 – “Genes overlap”**). The probability that an observed overlap between two drug/drug-state pairs was due to chance alone was evaluated with 100 rounds of Monte-Carlo simulation. More particularly, for each round a number equal to the number of genes participating in the statistically significant association rules for each drug/drug-state was randomly sampled from the total pool of genes participating in all the significant association rules (18216 genes). The random sampling was weighted by the frequency of occurrence of each gene in the sum of the significant association rules; hence a gene participating in numerous rules will have a greater probability of being picked in comparison to a gene participating in just a few rules. At the end of each round, the overlap between the randomly sampled gene-sets of the particular drug/drug-state pair under examination was recorded. At the end of the 100-round Monte-Carlo simulation, the distribution of the 100 measured overlaps (which was found to be normal by the Kolmogorov-Smirnov test for normality) was utilized to calculate the p-value of the actual overlap between the particular drug/drug-state pair (**Supplementary Table 6 – “p-values”**). This p-value represents the probability of the actual overlap to belong to the distribution of the randomly generated overlaps; hence the actual overlap being due to chance alone. If the actual overlap is located at the far right side of the random distribution the overlap is characterized as over-represented and statistically significant; hence non-randomly relevant. In contrast, if the actual overlap is located at the far left side of the random distribution the overlap is characterized as under-represented and statistically significant; hence non-randomly distant (scripts/script_measure_gene_overlaps_of_drugStates.R@GitHub).

### 6. Data Availability

All scripts, data objects, figures and tables have been deposited and can be accessed at the public GitHub repository (folder: “Vougas_DeepLearning”, https://github.com/kvougas/Vougas_DeepLearning)

## Author Contributions

K.V: study conception and design, scripting, bioinformatics analysis, experimental procedures, results interpretation, manuscript preparation and writing, T.J, A.P, A.A and M.K: scripting and data analysis, E.J, V.G, A.V, A.V and D.T: guidance and assistance in manuscript preparation, M.L, I.P, P.T: data interpretation and guidance, J.B: experimental procedures, results interpretation and guidance and V.G.G: study conception and design, experimental procedures, data analysis and interpretation, guidance and manuscript preparation. All authors discussed the results and commented on the manuscript

---

**Supplementary Figure 1: Relationships between metrics obtained through association rule mining. a)** Scatter plots presenting relation between confidence and support for 10,000 1-way rules based on top support, confidence and lift. **b)** Scatter plots presenting relation between confidence and support for 100,000 2-way rules based on top support, confidence and lift.

**Supplementary Figure 2: Silencing of *MAGI3*, *POF1B* and *PDIA3* genes in the cancer cell-lines (Supplementary Table 4). a)** Time line of experimental procedure. **b)** Cell viability assay of A549, H1299, MCF7 and Saos-2 cells after *POF1B*, *MAGI3* and *PDIA3* mRNA silencing. **c)** RT-PCR analysis of *POF1B*, *MAGI3*, *PDIA3* mRNA expression levels before and after RNA silencing in A549, H1299, MCF7 and Saos-2 cells.

**Supplementary Figure 3:** Distribution of cell-lines between (**a**) Training set and (**b**) Test set based on the tissue of origin. Original matrix was split upon blocked randomization (blocking factor: tissue type), where two thirds of the cell lines were randomly assigned to the Training sets and one third was assigned to the Test set.

**Drug resistance clustering based on the similarity of genes involved in the association rules with the resistance state of each drug.** Correlation of Jaccard-distance with rule confidence and lift for drug-to-drug association rules (**Supplementary Table 8 – “Res_res”**). Drugs are colour coded by drug type. Detailed information regarding the groups and the associated drug targets is located in **Supplementary Table 2**. Red clusters represent stable clusters.

**Supplementary Figure 5.** Randomness estimation of the degree of overlap between the drug-sensitivity and drug-resistance gene-sets for all drugs (Sensitive vs Resistant). Colour legend indicates significance of each overlap: *i)* blue – statistically significantly over-represented; *ii)* brown – statistically significantly under-represented, *iii)* white – random overlap. Detailed description of features corresponding to each rule-set, number of overlapping genes and p-values is reported in **Supplementary Table 6.**

**Supplementary Figure 6. Randomness estimation of the degree of overlap among the drug-sensitivity gene-sets and the drug-resistance gene-sets for all drugs (Sensitive vs Sensitive and Resistant vs Resistant).** Colour legend indicates significance of each overlap: *i)* blue – statistically significantly over-represented; *ii)* brown – statistically significantly under-represented, *iii)* white – random overlap. Detailed description of features corresponding to each rule-set, number of overlapping genes and p-values is reported in **Supplementary Table 6.**

**Supplementary Table 1**

**1- way rules,** Significant Rules (FDR<5%) from 1-way (A=>B) Association Rule Mining of the total dataset with minimum Support and Confidence levels at 0.3%. The Left Hand Side of the rules contain genes and tissue of origin while Right Hand Side Drug/Response pairs. GE= Gene Expression; CNV=Copy Number Variation

**2- way rules,** Significant Rules (FDR<5%) from 1-way (A+B=>C) Association Rule Mining of the total dataset with minimum Support and Confidence levels at 0.8%.

The Left Hand Side of the rules contain genes and tissue of origin while Right Hand Side Drug/Response pairs. GE= Gene Expression; CNV=Copy Number Variation

**NOTE:** The table legends for the remaining supplementary tables can be found in the respective Excel files. Supplementary table 1 due to extensive size could not be loaded in Excel, we therefore compiled a zip file containing two tab-delimited text files (1-way & 2-way rules).

## References

1. Halazonetis, T.D., Gorgoulis, V.G. & Bartek, J. An oncogene-induced DNA damage model for cancer development. Science 319, 1352–1355 (2008).

2. Galanos, P. et al. Chronic p53-independent p21 expression causes genomic instability by deregulating replication licensing. Nature cell biology 18, 777–789 (2016).

3. van't Veer, L.J. & Bernards, R. Enabling personalized cancer medicine through analysis of gene-expression patterns. Nature 452, 564–570 (2008).

4. Weinstein, J.N. Drug discovery: Cell lines battle cancer. Nature 483, 544–545 (2012).

5. Iorio, F. et al. A Landscape of Pharmacogenomic Interactions in Cancer. Cell 166, 740– 754 (2016).

6. Barretina, J. et al. The Cancer Cell Line Encyclopedia enables predictive modelling of anticancer drug sensitivity. Nature 483, 603–607 (2012).

7. Garnett, M.J. et al. Systematic identification of genomic markers of drug sensitivity in cancer cells. Nature 483, 570–575 (2012).

8. Shoemaker, R.H. The NCI60 human tumour cell line anticancer drug screen. Nature reviews. Cancer 6, 813–823 (2006).

9. Libbrecht, M.W. & Noble, W.S. Machine learning applications in genetics and genomics. Nature reviews. Genetics 16, 321–332 (2015).

10. Schmidhuber, J. Deep learning in neural networks: an overview. Neural networks : the official journal of the International Neural Network Society 61, 85–117 (2015).

11. LeCun, Y., Bengio, Y. & Hinton, G. Deep learning. Nature 521, 436–444 (2015).

12. Wang, C. et al. in Bioinformatics and Biomedicine (BIBM), 2014 IEEE International Conference on 67-70 (2014).

13. Xu, Y. et al. Deep Learning for Drug-Induced Liver Injury. Journal of chemical information and modeling 55, 2085–2093 (2015).

14. Aliper, A. et al. Deep Learning Applications for Predicting Pharmacological Properties of Drugs and Drug Repurposing Using Transcriptomic Data. Molecular pharmaceutics 13, 2524–2530 (2016).

15. Bengio, Y., Courville, A. & Vincent, P. Representation Learning: A Review and New Perspectives. IEEE transactions on pattern analysis and machine intelligence (2013).

16. Forbes, S.A. et al. COSMIC: exploring the world's knowledge of somatic mutations in human cancer. Nucleic acids research 43, D805–811 (2015).

17. Costello, J.C. et al. A community effort to assess and improve drug sensitivity prediction algorithms. Nature biotechnology 32, 1202–1212 (2014).

18. Masica, D.L. & Karchin, R. Collections of simultaneously altered genes as biomarkers of cancer cell drug response. Cancer research 73, 1699–1708 (2013).

19. Tan, P.-N., Kumar, V. & Srivastava, J. Selecting the right objective measure for association analysis. Information Systems 29, 293–313 (2004).

20. Agrawal, R., Imieli, T., #324, ski & Swami, A. in Proceedings of the 1993 ACM SIGMOD international conference on Management of data 207–216 (ACM, Washington, D.C., USA; 1993).

21. Kelland, L.R., Sharp, S.Y., Rogers, P.M., Myers, T.G. & Workman, P. DT-Diaphorase expression and tumor cell sensitivity to 17-allylamino, 17-demethoxygeldanamycin, an inhibitor of heat shock protein 90. Journal of the National Cancer Institute 91, 1940– 1949 (1999).

22. Muller, C.R. et al. Potential for treatment of liposarcomas with the MDM2 antagonist Nutlin-3A. International journal of cancer 121, 199–205 (2007).

23. Rusnak, D.W. et al. Assessment of epidermal growth factor receptor (EGFR, ErbB1) and HER2 (ErbB2) protein expression levels and response to lapatinib (Tykerb, GW572016) in an expanded panel of human normal and tumour cell lines. Cell proliferation 40, 580–594 (2007).

24. Long, G.V. et al. Combined BRAF and MEK inhibition versus BRAF inhibition alone in melanoma. The New England journal of medicine 371, 1877–1888 (2014).

25. Carracedo, A. & Pandolfi, P.P. The PTEN-PI3K pathway: of feedbacks and cross-talks. Oncogene 27, 5527–5541 (2008).

26. Rodriguez-Escudero, I. et al. A comprehensive functional analysis of PTEN mutations: implications in tumor- and autism-related syndromes. Human molecular genetics 20, 4132–4142 (2011).

27. Knudson, A.G., Jr. Mutation and cancer: statistical study of retinoblastoma. Proceedings of the National Academy of Sciences of the United States of America 68, 820–823 (1971).

28. Eisfeld, A.K. et al. Mutational Landscape and Gene Expression Patterns in Adult Acute Myeloid Leukemias with Monosomy 7 as a Sole Abnormality. Cancer research 77, 207– 218 (2017).

29. Jin, X. et al. The ID1-CULLIN3 Axis Regulates Intracellular SHH and WNT Signaling in Glioblastoma Stem Cells. Cell reports 16, 1629–1641 (2016).

30. Li, W. et al. An essential role for the Id1/PI3K/Akt/NFkB/survivin signalling pathway in promoting the proliferation of endothelial progenitor cells in vitro. Molecular and cellular biochemistry 363, 135–145 (2012).

31. Tominaga, K. et al. Addiction to the IGF2-ID1-IGF2 circuit for maintenance of the breastcancer stem-like cells. Oncogene (2016).

32. Murase, R. et al. Suppression of invasion and metastasis in aggressive salivary cancer cells through targeted inhibition of ID1 gene expression. Cancer letters 377, 11–16 (2016).

33. Fransecky, L., Mochmann, L.H. & Baldus, C.D. Outlook on PI3K/AKT/mTOR inhibition in acute leukemia. Molecular and cellular therapies 3, 2 (2015).

34. King, M.A., Ganley, I.G. & Flemington, V. Inhibition of cholesterol metabolism underlies synergy between mTOR pathway inhibition and chloroquine in bladder cancer cells. Oncogene 35, 4518–4528 (2016).

35. Meng, Y.S., Khoury, H., Dick, J.E. & Minden, M.D. Oncogenic potential of the transcription factor LYL1 in acute myeloblastic leukemia. Leukemia 19, 1941–1947 (2005).

36. Zhong, Y., Jiang, L., Hiai, H., Toyokuni, S. & Yamada, Y. Overexpression of a transcription factor LYL1 induces T- and B-cell lymphoma in mice. Oncogene 26, 6937– 6947 (2007).

37. Bertacchini, J. et al. Targeting PI3K/AKT/mTOR network for treatment of leukemia. Cellular and molecular life sciences : CMLS 72, 2337–2347 (2015).

38. Blachly, J.S. & Baiocchi, R.A. Targeting PI3-kinase (PI3K), AKT and mTOR axis in lymphoma. British journal of haematology 167, 19–32 (2014).

39. Noll, J.E. et al. SAMSN1 is a tumor suppressor gene in multiple myeloma. Neoplasia 16, 572–585 (2014).

40. Kanda, M. et al. Prognostic relevance of SAMSN1 expression in gastric cancer. Oncology letters 12, 4708–4716 (2016).

41. Yamada, H. et al. Detailed characterization of a homozygously deleted region corresponding to a candidate tumor suppressor locus at 21q11-21 in human lung cancer. Genes, chromosomes & cancer 47, 810–818 (2008).

42. Sueoka, S. et al. Suppression of SAMSN1 Expression is Associated with the Malignant Phenotype of Hepatocellular Carcinoma. Annals of surgical oncology 22 Suppl 3, S1453–1460 (2015).

43. Yan, Y. et al. SAMSN1 is highly expressed and associated with a poor survival in glioblastoma multiforme. PloS one 8, e81905 (2013).

44. Haar, C.P. et al. Drug resistance in glioblastoma: a mini review. Neurochemical research 37, 1192–1200 (2012).

45. Sami, A. & Karsy, M. Targeting the PI3K/AKT/mTOR signaling pathway in glioblastoma: novel therapeutic agents and advances in understanding. Tumour biology : the journal of the International Society for Oncodevelopmental Biology and Medicine 34, 1991–2002 (2013).

46. Planchard, D. & Le Pechoux, C. Small cell lung cancer: new clinical recommendations and current status of biomarker assessment. European journal of cancer 47 Suppl 3, S272–283 (2011).

47. Yeh, J.J. et al. Comparison of chemotherapy response with P-glycoprotein, multidrug resistance-related protein-1, and lung resistance-related protein expression in untreated small cell lung cancer. Lung 183, 177–183 (2005).

48. Kiaris, H., Schally, A.V. & Varga, J.L. Suppression of tumor growth by growth hormone-releasing hormone antagonist JV-1-36 does not involve the inhibition of autocrine production of insulin-like growth factor II in H-69 small cell lung carcinoma. Cancer letters 161, 149–155 (2000).

49. Perez, R. et al. Antagonistic analogs of growth hormone-releasing hormone increase the efficacy of treatment of triple negative breast cancer in nude mice with doxorubicin; A preclinical study. Oncoscience 1, 665–673 (2014).

50. Hohla, F. et al. Synergistic inhibition of growth of lung carcinomas by antagonists of growth hormone-releasing hormone in combination with docetaxel. Proceedings of the National Academy of Sciences of the United States of America 103, 14513–14518 (2006).

51. Yang, Y.A., Zhang, G.M., Feigenbaum, L. & Zhang, Y.E. Smad3 reduces susceptibility to hepatocarcinoma by sensitizing hepatocytes to apoptosis through downregulation of Bcl-2. Cancer cell 9, 445–457 (2006).

52. Wu, Y. et al. Interaction of the tumor suppressor PTEN/MMAC with a PDZ domain of MAGI3, a novel membrane-associated guanylate kinase. The Journal of biological chemistry 275, 21477–21485 (2000).

53. Pal, I. & Mandal, M. PI3K and Akt as molecular targets for cancer therapy: current clinical outcomes. Acta pharmacologica Sinica 33, 1441–1458 (2012).

54. Strickland, S., Wasserman, J.K., Giassi, A., Djordjevic, B. & Parra-Herran, C. Immunohistochemistry in the Diagnosis of Mucinous Neoplasms Involving the Ovary: The Added Value of SATB2 and Biomarker Discovery Through Protein Expression Database Mining. International journal of gynecological pathology : official journal of the International Society of Gynecological Pathologists 35, 191–208 (2016).

55. Padovano, V. et al. The POF1B candidate gene for premature ovarian failure regulates epithelial polarity. Journal of cell science 124, 3356–3368 (2011).

56. Takeshita, H. et al. Actin organization associated with the expression of multidrug resistant phenotype in osteosarcoma cells and the effect of actin depolymerization on drug resistance. Cancer letters 126, 75–81 (1998).

57. Ramirez-Rangel, I. et al. Regulation of mTORC1 complex assembly and signaling by GRp58/ERp57. Molecular and cellular biology 31, 1657–1671 (2011).

58. Hussmann, M. et al. Depletion of the thiol oxidoreductase ERp57 in tumor cells inhibits proliferation and increases sensitivity to ionizing radiation and chemotherapeutics. Oncotarget 6, 39247–39261 (2015).

59. Breiman, L. Bagging Predictors. Machine Learning 24, 123–140 (1996).

60. Srivastava, N., Hinton, G., Krizhevsky, A., Sutskever, I. & Salakhutdinov, R. Dropout: a simple way to prevent neural networks from overfitting. J. Mach. Learn. Res. 15, 1929– 1958 (2014).

61. Zou, H. & Hastie, T. Regularization and variable selection via the elastic net. Journal of the Royal Statistical Society: Series B (Statistical Methodology) 67, 301–320 (2005).

62. Chen, X. & Ishwaran, H. Random forests for genomic data analysis. Genomics 99, 323– 329 (2012).

63. Gonen, M. & Margolin, A.A. Drug susceptibility prediction against a panel of drugs using kernelized Bayesian multitask learning. Bioinformatics 30, i556–563 (2014).

64. Metz, C.E. Basic principles of ROC analysis. Seminars in nuclear medicine 8, 283–298 (1978).

65. Youden, W.J. Index for rating diagnostic tests. Cancer 3, 32–35 (1950).

66. Matthews, B.W. Comparison of the predicted and observed secondary structure of T4 phage lysozyme. Biochimica et biophysica acta 405, 442–451 (1975).

67. Craft, D., Ferranti, D. & Krane, D. The value of prior knowledge in machine learning of complex network systems. bioRxiv (2016).

68. Breiman, L. Random Forests. Machine Learning 45, 5–32 (2001).

69. Chen, D., Liu, X., Yang, Y., Yang, H. & Lu, P. Systematic synergy modeling: understanding drug synergy from a systems biology perspective. BMC systems biology 9, 56 (2015).

70. Burotto, M., Chiou, V.L., Lee, J.M. & Kohn, E.C. The MAPK pathway across different malignancies: a new perspective. Cancer 120, 3446–3456 (2014).

71. Jia, J. et al. Mechanisms of drug combinations: interaction and network perspectives. Nature reviews. Drug discovery 8, 111–128 (2009).

72. Shi, F. et al. Ribavirin Inhibits the Activity of mTOR/eIF4E, ERK/Mnk1/eIF4E Signaling Pathway and Synergizes with Tyrosine Kinase Inhibitor Imatinib to Impair Bcr-Abl Mediated Proliferation and Apoptosis in Ph+ Leukemia. PloS one 10, e0136746 (2015).

73. Badura, S. et al. Differential effects of selective inhibitors targeting the PI3K/AKT/mTOR pathway in acute lymphoblastic leukemia. PloS one 8, e80070 (2013).

74. Miura, K. et al. The combination of olaparib and camptothecin for effective radiosensitization. Radiation oncology 7, 62 (2012).

75. Minami, D. et al. Synergistic effect of olaparib with combination of cisplatin on PTEN-deficient lung cancer cells. Molecular cancer research : MCR 11, 140–148 (2013).

76. Tahara, M. et al. The use of Olaparib (AZD2281) potentiates SN-38 cytotoxicity in colon cancer cells by indirect inhibition of Rad51-mediated repair of DNA double-strand breaks. Molecular cancer therapeutics 13, 1170–1180 (2014).

77. Prahallad, A. et al. Unresponsiveness of colon cancer to BRAF(V600E) inhibition through feedback activation of EGFR. Nature 483, 100–103 (2012).

78. Corcoran, R.B. et al. EGFR-mediated re-activation of MAPK signaling contributes to insensitivity of BRAF mutant colorectal cancers to RAF inhibition with vemurafenib. Cancer discovery 2, 227–235 (2012).

79. Uitdehaag, J.C. et al. Selective Targeting of CTNBB1-, KRAS- or MYC-Driven Cell Growth by Combinations of Existing Drugs. PloS one 10, e0125021 (2015).

80. Haagensen, E.J., Kyle, S., Beale, G.S., Maxwell, R.J. & Newell, D.R. The synergistic interaction of MEK and PI3K inhibitors is modulated by mTOR inhibition. British journal of cancer 106, 1386–1394 (2012).

81. Place, R.F., Noonan, E.J. & Giardina, C. HDAC inhibition prevents NF-kappa B activation by suppressing proteasome activity: down-regulation of proteasome subunit expression stabilizes I kappa B alpha. Biochemical pharmacology 70, 394–406 (2005).

82. Kaler, P., Sasazuki, T., Shirasawa, S., Augenlicht, L. & Klampfer, L. HDAC2 deficiency sensitizes colon cancer cells to TNFalpha-induced apoptosis through inhibition of NF-kappaB activity. Experimental cell research 314, 1507–1518 (2008).

83. Imre, G., Gekeler, V., Leja, A., Beckers, T. & Boehm, M. Histone deacetylase inhibitors suppress the inducibility of nuclear factor-kappaB by tumor necrosis factor-alpha receptor-1 down-regulation. Cancer research 66, 5409–5418 (2006).

84. Orlikova, B. et al. Natural chalcones as dual inhibitors of HDACs and NF-kappaB. Oncology reports 28, 797–805 (2012).

85. Dhanyamraju, P.K., Holz, P.S., Finkernagel, F., Fendrich, V. & Lauth, M. Histone deacetylase 6 represents a novel drug target in the oncogenic Hedgehog signaling pathway. Molecular cancer therapeutics 14, 727–739 (2015).

86. Zhao, J., Quan, H., Xie, C. & Lou, L. NL-103, a novel dual-targeted inhibitor of histone deacetylases and hedgehog pathway, effectively overcomes vismodegib resistance conferred by Smo mutations. Pharmacology research & perspectives 2, e00043 (2014).

87. Lo, R.S. Receptor tyrosine kinases in cancer escape from BRAF inhibitors. Cell research 22, 945–947 (2012).

88. Deng, Z.J. et al. Layer-by-layer nanoparticles for systemic codelivery of an anticancer drug and siRNA for potential triple-negative breast cancer treatment. ACS nano 7, 9571– 9584 (2013).

89. Hornik, K., Stinchcombe, M. & White, H. Multilayer feedforward networks are universal approximators. Neural Networks 2, 359–366 (1989).

90. Jiang, G. et al. In human leukemia cells ephrin-B-induced invasive activity is supported by Lck and is associated with reassembling of lipid raft signaling complexes. Molecular cancer research : MCR 6, 291–305 (2008).

91. Blair, L.J., Zhang, B. & Dickey, C.A. Potential synergy between tau aggregation inhibitors and tau chaperone modulators. Alzheimer's research & therapy 5, 41 (2013).

92. Zhang, Q. et al. P-glycoprotein-evading anti-tumor activity of a novel tubulin and HSP90 dual inhibitor in a non-small-cell lung cancer model. Journal of pharmacological sciences 126, 66–76 (2014).

93. Das, V. et al. Pathophysiologically relevant in vitro tumor models for drug screening. Drug discovery today 20, 848–855 (2015).

94. Siolas, D. & Hannon, G.J. Patient-derived tumor xenografts: transforming clinical samples into mouse models. Cancer research 73, 5315–5319 (2013).

95. R Core Team (R Foundation for Statistical Computing, Vienna, Austria; 2016).

96. Hahsler, M. & Karpienko, R. Visualizing association rules in hierarchical groups. Journal of Business Economics, 1–19 (2016).

97. Eliades, T., Pratsinis, H., Athanasiou, A.E., Eliades, G. & Kletsas, D. Cytotoxicity and estrogenicity of Invisalign appliances. American journal of orthodontics and dentofacial orthopedics : official publication of the American Association of Orthodontists, its constituent societies, and the American Board of Orthodontics 136, 100–103 (2009).

98. Sing, T., Sander, O., Beerenwinkel, N. & Lengauer, T. ROCR: visualizing classifier performance in R. Bioinformatics 21, 3940–3941 (2005).

99. Oshiro, T.M., Perez, P.S. & Baranauskas, J.A. in Machine Learning and Data Mining in Pattern Recognition. (ed. P. Petra) (Springer-Verlag Berlin Heidelberg, 2012).

100. Menden, M.P. et al. Machine learning prediction of cancer cell sensitivity to drugs based on genomic and chemical properties. PloS one 8, e61318 (2013).

101. Jaccard, P. THE DISTRIBUTION OF THE FLORA IN THE ALPINE ZONE.1. New Phytologist 11, 37–50 (1912).

102. Oksanen, J. et al. vegan: Community Ecology Package. (2016).

103. Suzuki, R. & Shimodaira, H. Pvclust: an R package for assessing the uncertainty in hierarchical clustering. Bioinformatics 22, 1540–1542 (2006).

104. Bostock, M., Ogievetsky, V. & Heer, J. D3: Data-Driven Documents. IEEE Trans. Visualization \& Comp. Graphics (Proc. InfoVis) (2011).

