## Supplementary Materials for "Deep Learning and Association Rule Mining for Predicting Drug Response in Cancer. A Personalised Medicine Approach"

### Slide 1
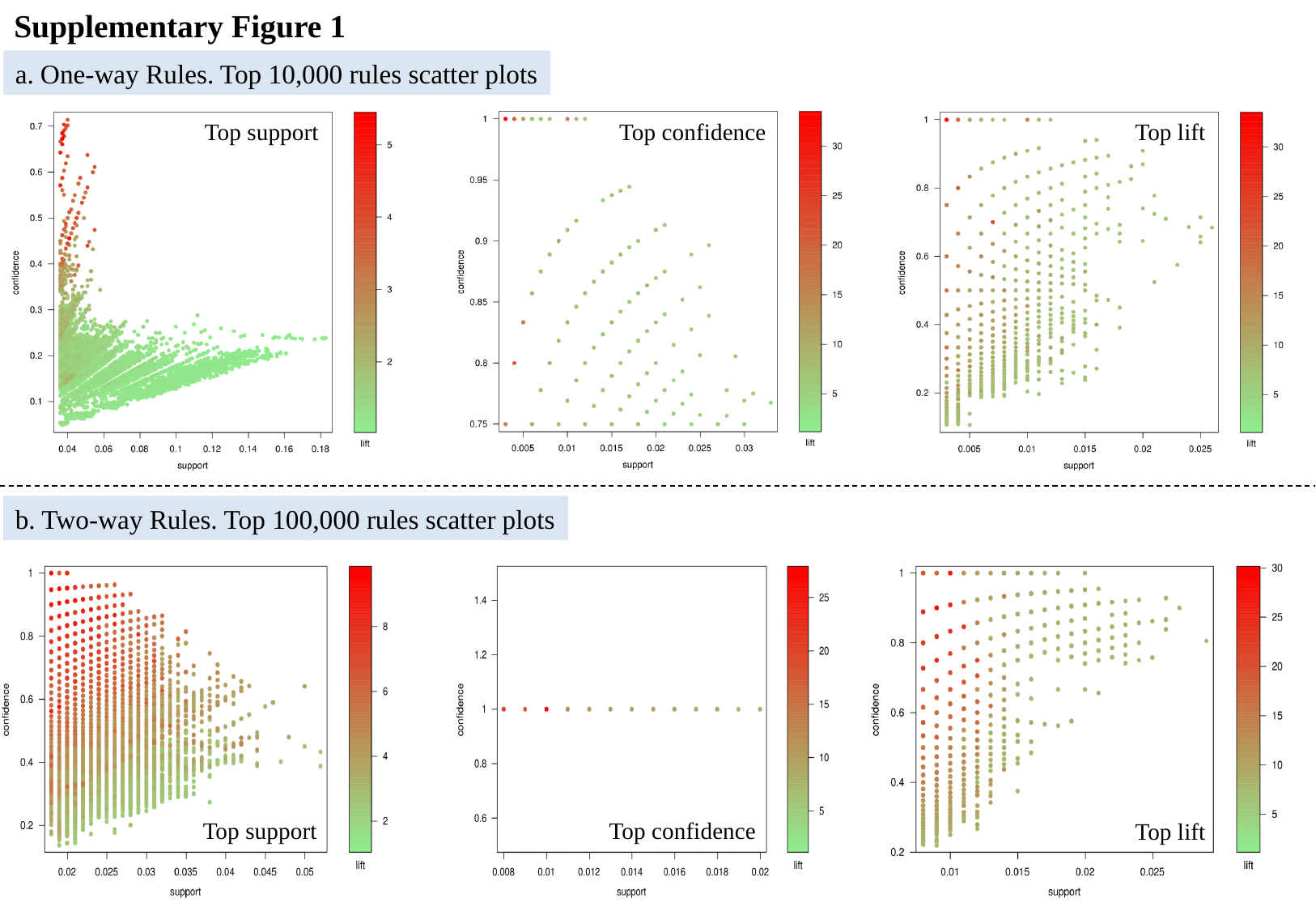

Supplementary Figure 1
a. One-way Rules. Top 10,000 rules scatter plots
Top support
Top confidence
Top lift
b. Two-way Rules. Top 100,000 rules scatter plots
Top support
Top confidence
Top lift
