## Supplementary Materials for "Deep Learning and Association Rule Mining for Predicting Drug Response in Cancer. A Personalised Medicine Approach"

### Slide 1
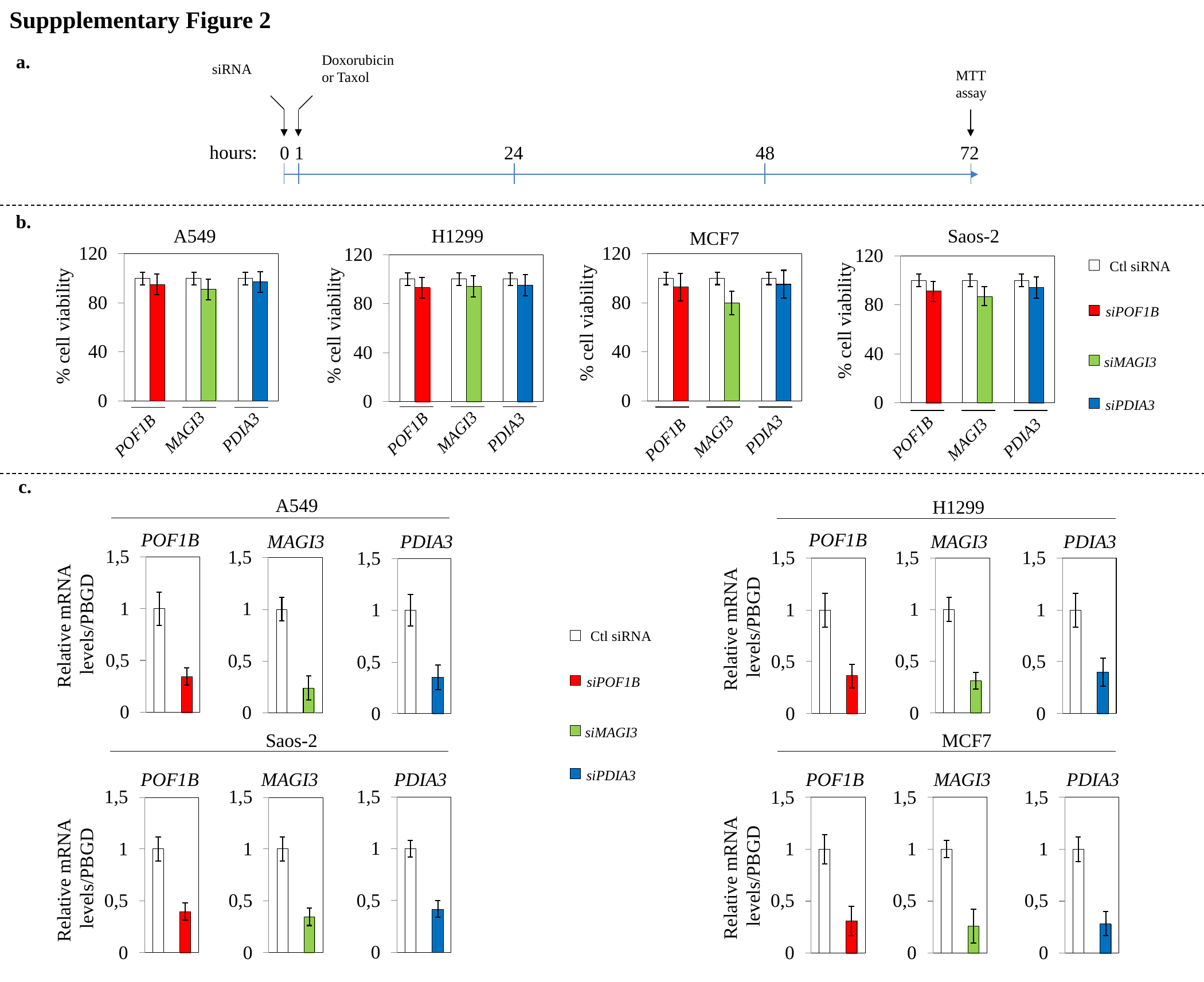

Suppplementary Figure 2
a.
Doxorubicin
or Taxol
siRNA
MTT
assay
hours:
0
1
24
48
72
b.
A549
% cell viability
MAGI3
PDIA3
POF1B
H1299
% cell viability
MAGI3
PDIA3
POF1B
Saos-2
% cell viability
MAGI3
PDIA3
POF1B
MCF7
% cell viability
MAGI3
PDIA3
POF1B
Ctl siRNA
siPOF1B
siMAGI3
siPDIA3
c.
A549
POF1B
MAGI3
PDIA3
Relative mRNA
levels/PBGD
H1299
POF1B
MAGI3
PDIA3
Relative mRNA
levels/PBGD
Ctl siRNA
siPOF1B
siMAGI3
MCF7
POF1B
MAGI3
PDIA3
Relative mRNA
levels/PBGD
Saos-2
POF1B
MAGI3
PDIA3
Relative mRNA
levels/PBGD
siPDIA3
