## Supplementary figures and images for "Deep Learning and Association Rule Mining for Predicting Drug Response in Cancer. A Personalised Medicine Approach"

### Supplementary Materials

#### Slide 1
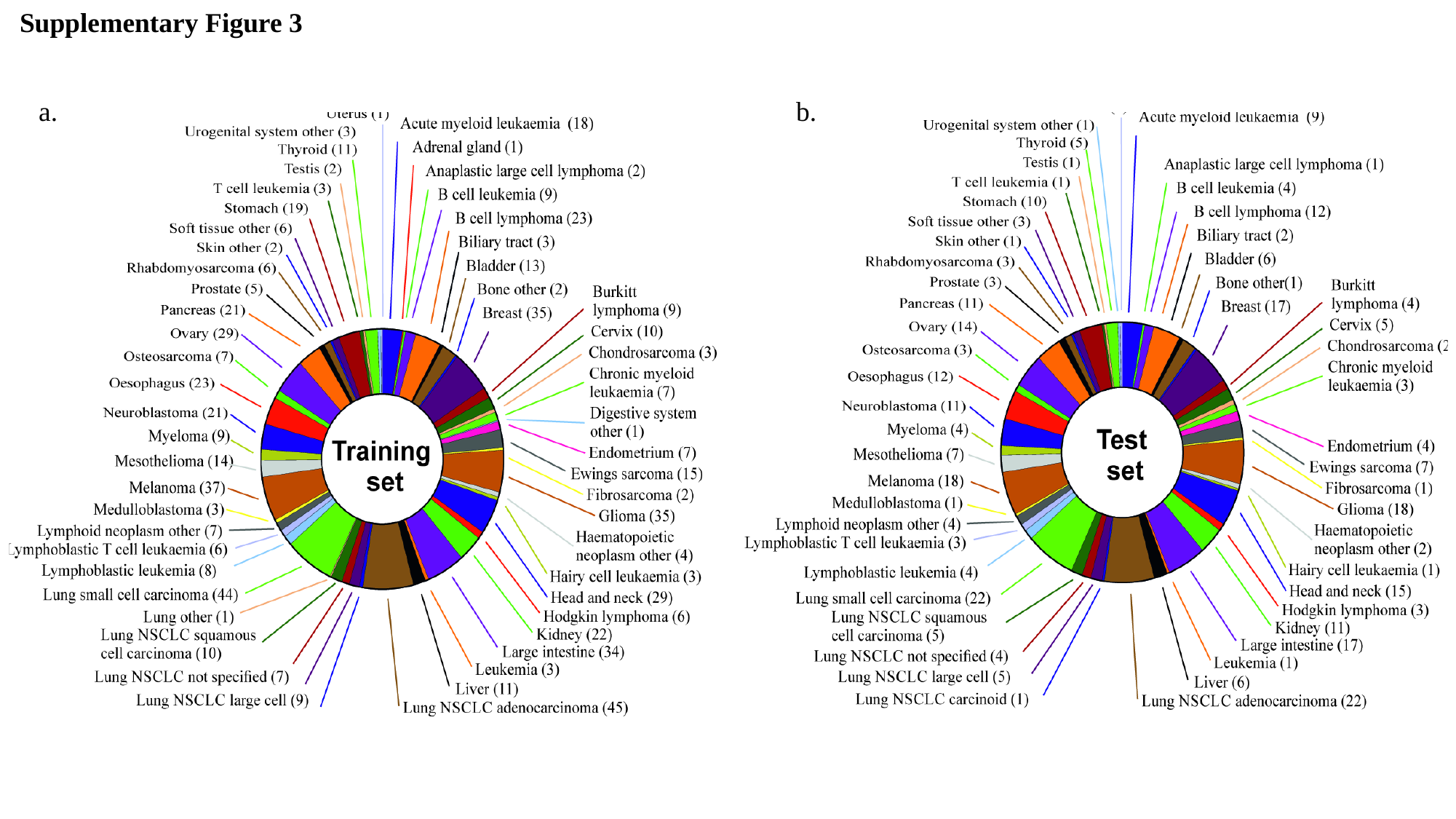

Supplementary Figure 3
a.
b.

### Supplementary Materials

#### Slide 1
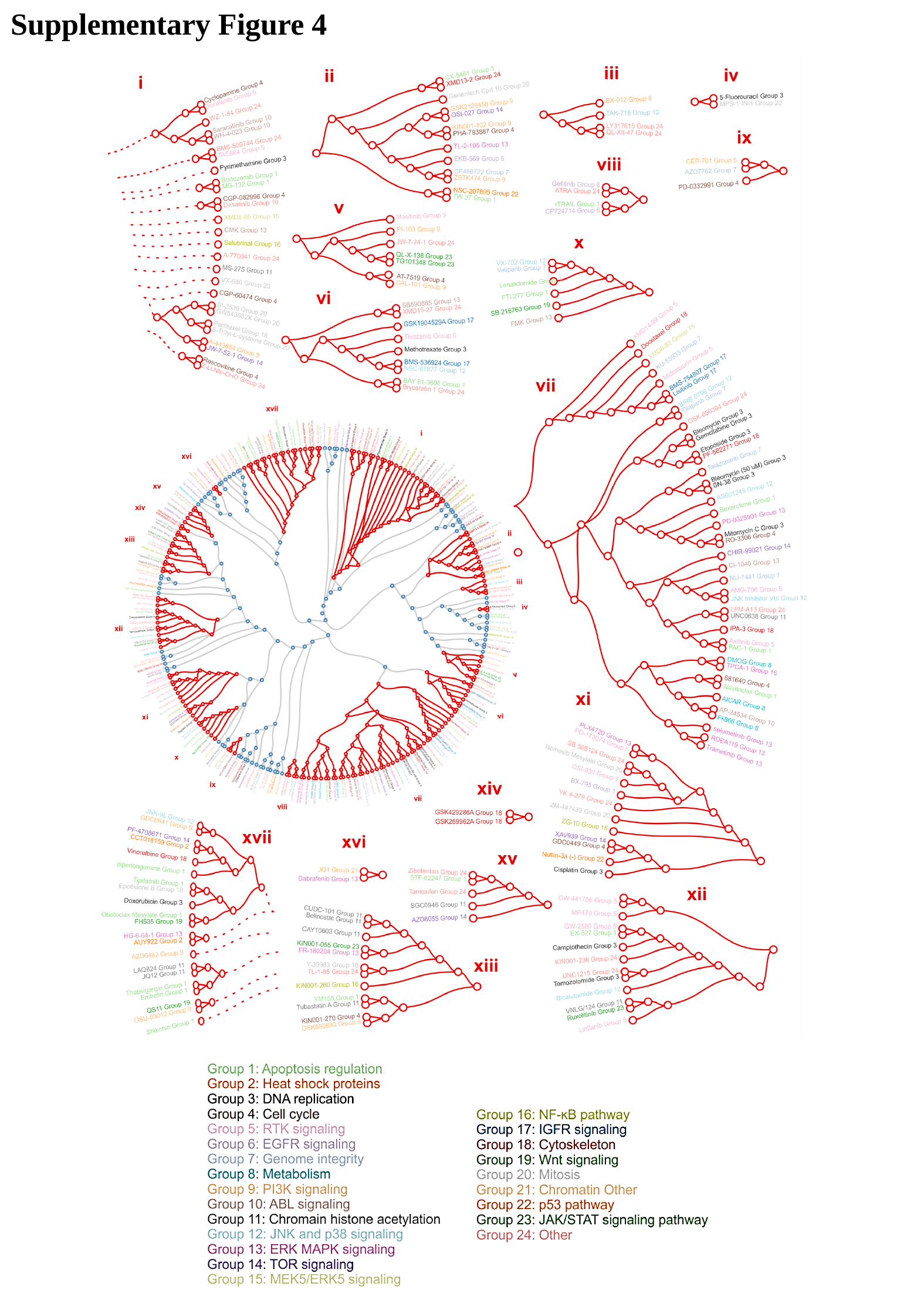

Supplementary Figure 4
