## Supplementary Materials for "Deep Learning and Association Rule Mining for Predicting Drug Response in Cancer. A Personalised Medicine Approach"

### Slide 1
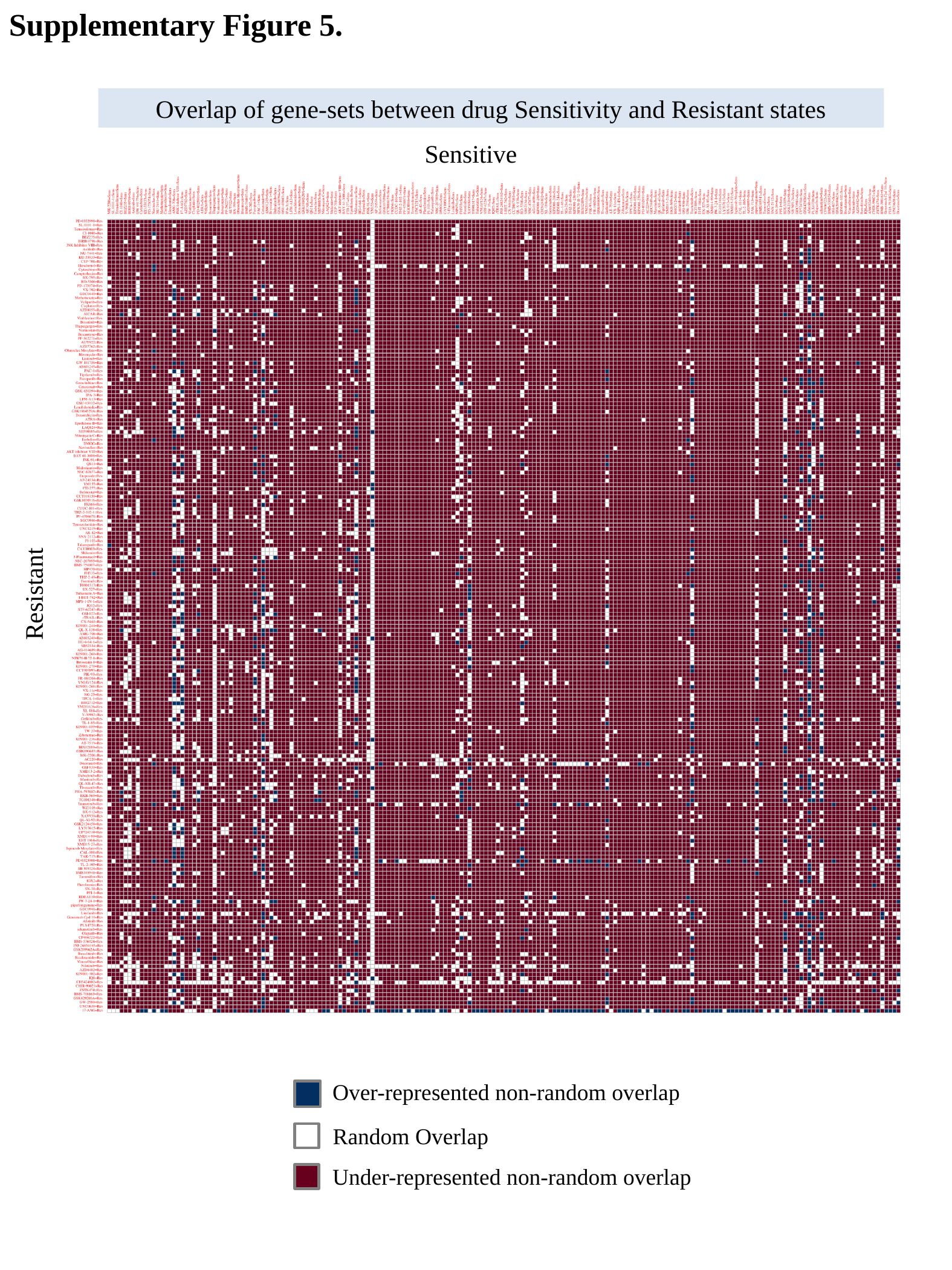

Supplementary Figure 5.
Overlap of gene-sets between drug Sensitivity and Resistant states
Sensitive
Resistant
Over-represented non-random overlap
Random Overlap
Under-represented non-random overlap
