## Supplementary Materials for "Deep Learning and Association Rule Mining for Predicting Drug Response in Cancer. A Personalised Medicine Approach"

### Slide 1
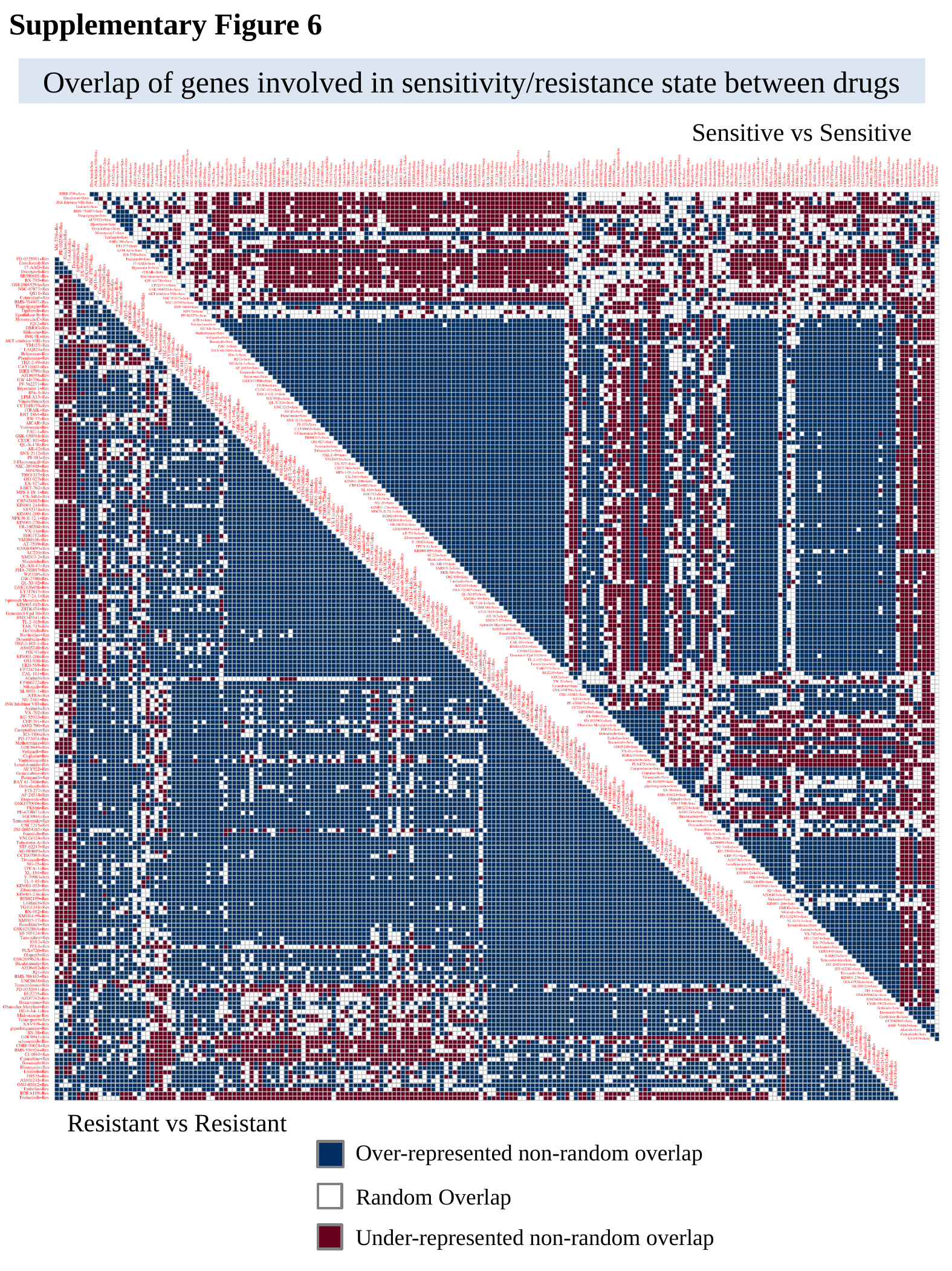

Supplementary Figure 6
Overlap of genes involved in sensitivity/resistance state between drugs
Sensitive vs Sensitive
Resistant vs Resistant
Over-represented non-random overlap
Random Overlap
Under-represented non-random overlap
